# Engineering the effector specificity of the macrolide-sensing transcription factor MphR for the detection of sesquiterpenes

**DOI:** 10.64898/2026.10.16.759199

**Authors:** Cassidy L. Oliverio, Alissa M. Meo, Stefanie Chen, Gavin J. Williams

## Abstract

Terpenes are a large, diverse class of natural products used as flavors, fragrances, pharmaceuticals, and more, yet obtaining them often requires either large quantities of biomass or complex chemical reactions. Directed evolution of key enzymes and pathways could improve biosynthetic yields, but most sesquiterpenes lack high-throughput screening methods to search large libraries of variants. To alleviate this bottleneck, we developed the first reported genetically encoded biosensor to detect four highly valuable sesquiterpenes: valencene, nootkatone, artemisinin, and cedrol. Starting with the macrolide transcription factor biosensor MphR, we screened a panel of mutants against the four sesquiterpenes, including MphR-13 (T17N/V18I/K21L/S106L/H147L/P161L), which readily detected all four sesquiterpenes. Computational modeling showed that the mutations reduced the size of the ligand-binding pocket compared with the wild type, which positively correlated with the biosensor’s fluorescence output. Additionally, swapping the fluorescence reporter improved the sensor’s response 15-20-fold, yielding a generalist sesquiterpene biosensor that can be further developed and applied to the directed evolution of sesquiterpene biosynthesis.

**Graphical Abstract:** 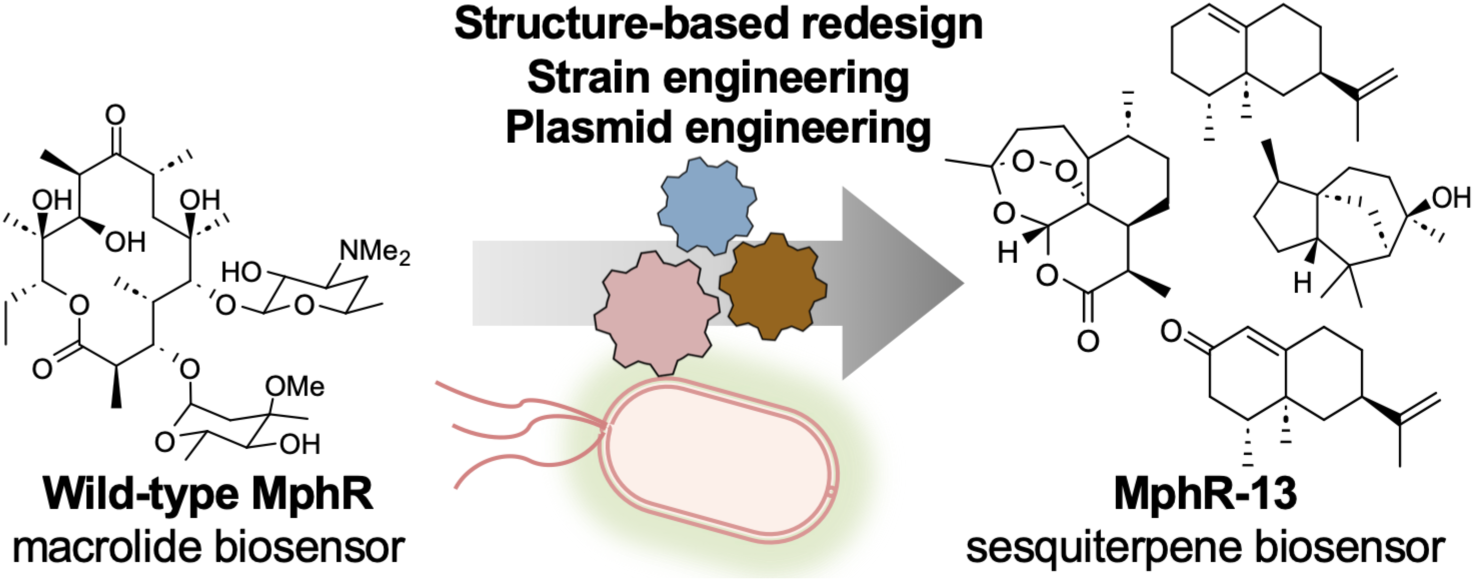

## 1. INTRODUCTION

Terpenes are a large class of secondary metabolites with applications across various fields that would benefit from high-throughput screening to improve accessibility. Terpenes are commonly used in various industries as flavors (e.g., valencene, myrcene), fragrances (e.g., geraniol, nootkatone), and materials (e.g., natural rubber), as well as in medicine for anti-cancer (e.g., Taxol) and anti-malaria (e.g., artemisinin) treatments [1]. In nature, terpenes are universally derived from the 5-carbon building blocks isopentenyl pyrophosphate (IPP) and dimethylallyl diphosphate (DMAPP) [1]. These building blocks are subsequently polymerized into a variety of chain lengths, including 15-carbon prenyl pyrophosphates that yield sesquiterpenes, such as artemisinin and nootkatone. Following cyclization, various enzymes further decorate the terpene scaffolds to generate the final product. For commercial and pharmaceutical purposes, the main method for producing many sesquiterpenes remains extraction from natural sources, despite their presence in minute quantities [2,3]. Many of these terpenes can be made via traditional synthetic chemistry but often use heavy-metal catalysts and result in low product titers [4–6].

Microbial production platforms offer a potentially sustainable alternative to plant extraction and chemical synthesis. However, microbial titers have not yet reached levels high enough to justify abandoning plant extraction. For high-level production, competition from native terpene pathways and various rate-limiting steps in the biosynthetic pathway impact final titers, and even the best yields fall short of industry demand [7–12]. Rational optimization of terpene biosynthetic pathways to overcome bottlenecks is challenging and often requires assaying hundreds of strains, pathways, and protein variants. Large combinatorial libraries of pathway variants or mutant enzyme libraries could be explored to rapidly identify and overcome simultaneous bottlenecks [13–15]. Critically, however, this approach requires efficient screening or selection to identify variants that yield higher product titers.

Engineered, genetically encoded biosensors based on allosteric transcription factors are valuable tools for screening pathway and enzyme libraries [16,17]. In the absence of a cognate effector, the transcription factor binds to the operator sequence upstream of a reporter gene, such as Green Fluorescent Protein (GFP). In the presence of the effector, its binding to the transcription factor induces a conformational change and dissociation from the operator sequence, leading to reporter expression and subsequent quantification [16]. Transcription factor biosensors have been utilized to guide the optimization of various natural product biosynthetic pathways, including those for alkaloids and pyrones [15,18]. However, for terpenes, most transcription factor biosensors have been restricted to monoterpenes (C10), including β-myrcene, camphor, various bicyclic terpenes, α-terpineol, and perillyl alcohol [19–22]. As an illustration of their utility, the MyrR biosensor was used to identify strains with 10-fold higher β-myrcene production [19]. Additionally, sensors have been developed for larger steroidal detecting sensors (C30) [23]. However, transcription factor biosensors for C15 terpenes have not yet been described.

Given the lack of known transcription factors that respond to C15 and larger terpenes, and inspired by previous examples [17,22] of engineering transcription factor effector specificity, we explored whether transcription factors with known promiscuity toward unrelated compounds could serve as starting points for directed evolution toward C15 terpenes. Notably, we previously used directed evolution to expand the effector specificity of the promiscuous macrolide-sensing transcription factor MphR [24–26] (**Fig. 1A**) to include macrolides that the wild-type biosensor poorly detects and 12/14-membered macrolactones that lack the sugars and phosphate required by the wild-type biosensor [27–29]. Given that the 14-membered macrolactone core resembles the C15-scaffolds of sesquiterpenes in terms of size and overall composition, we hypothesized that MphR may be a starting point for the development of transcription factor biosensors for the detection of sesquiterpenes.

**Figure 1.**
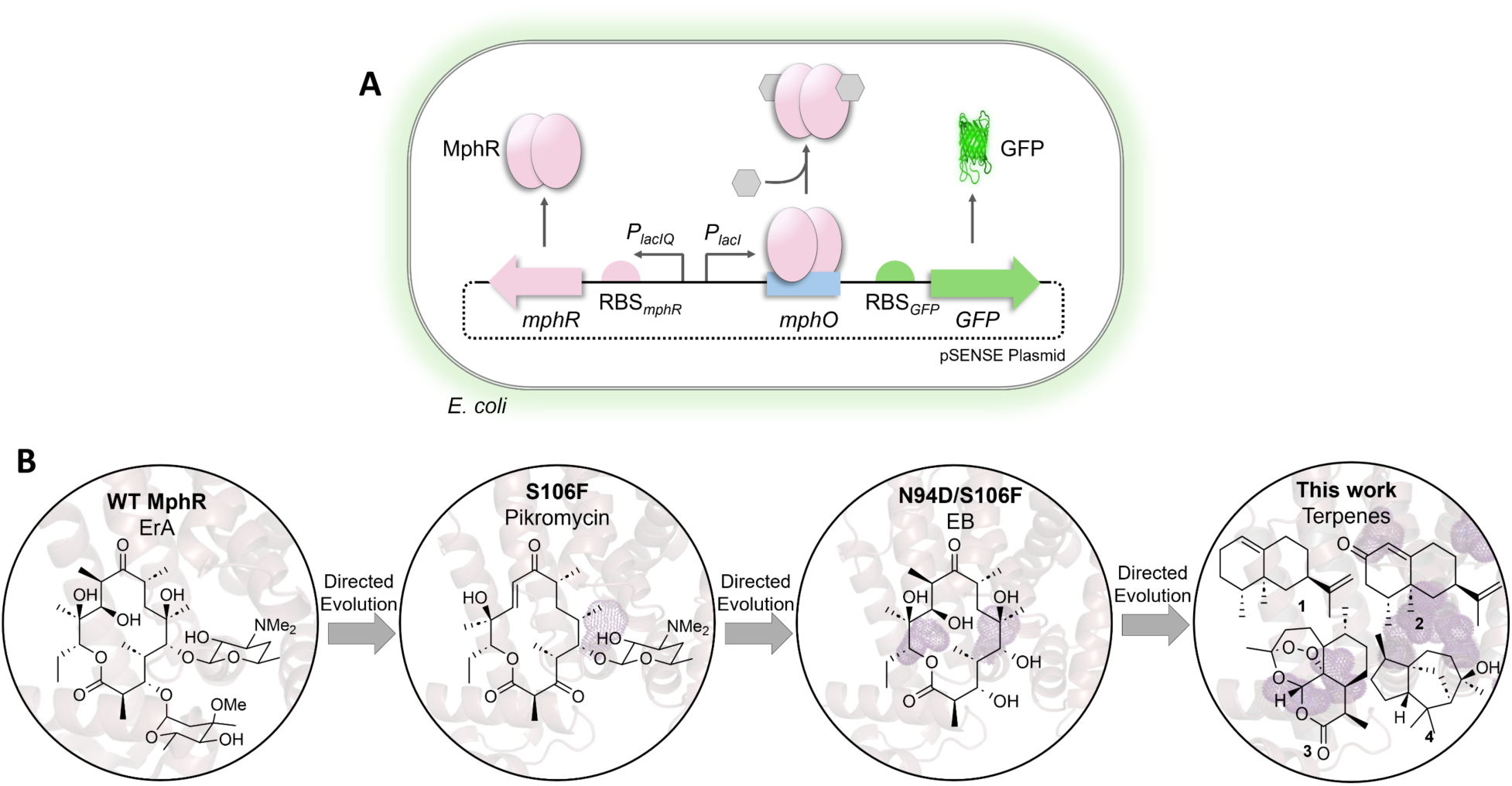
**A**) The MphR biosensor gene circuit. The presence of the ligand (grayed hexagon) triggers release of the transcription factor protein from the DNA, allowing for GFP expression. **B**) The directed evolution path of MphR, progressing via the native MphR ligand, erythromycin A (ErA), pikromycin (Pik), and the aglycone erythronolide B (EB) as key intermediates. This work focuses on detecting the sesquiterpenes valencene (**1**), nootkatone (**2**), artemisinin (**3**), and cedrol (**4**).

Here, we show that MphR can serve as a platform for developing effective sesquiterpene biosensors (**Fig. 1B**). A systematic campaign of screening current in-house MphR variants and subsequent whole-plasmid optimization resulted in the variant αMphR-13, which has a response ∼15-fold higher than that of the wild-type MphR toward the sesquiterpenes (+)-valencene (**1**), (+)-nootkatone (**2**), (+)-artemisinin (**3**), and (+)-cedrol (**4**). These results demonstrate that the effector specificity of MphR is highly evolvable and that early products of its directed evolution are highly promiscuous, being de-repressed by polyketides and terpenes. The engineered MphR variants set the stage for developing high-throughput approaches to guide sesquiterpene pathway engineering.

## 2. RESULTS AND DISCUSSION

### 2.1. Assembly of an MphR Variant Library and Sesquiterpene Panel

We assembled a small sesquiterpene panel for initial screening of MphR variants based on their valuable applications [30–33], suboptimal heterologous titers [7,9,10,34], and their representation of diverse skeletal structures and functionalities. The panel includes the hydrocarbon valencene (**1**), the ketone (+)-nootkatone (**2**), the highly oxidized multi-ring scaffold of (+)-artemisinin (**3**), and the tricyclic alcohol (+)-cedrol (**4**) (**Fig. 2**).

**Figure 2.**
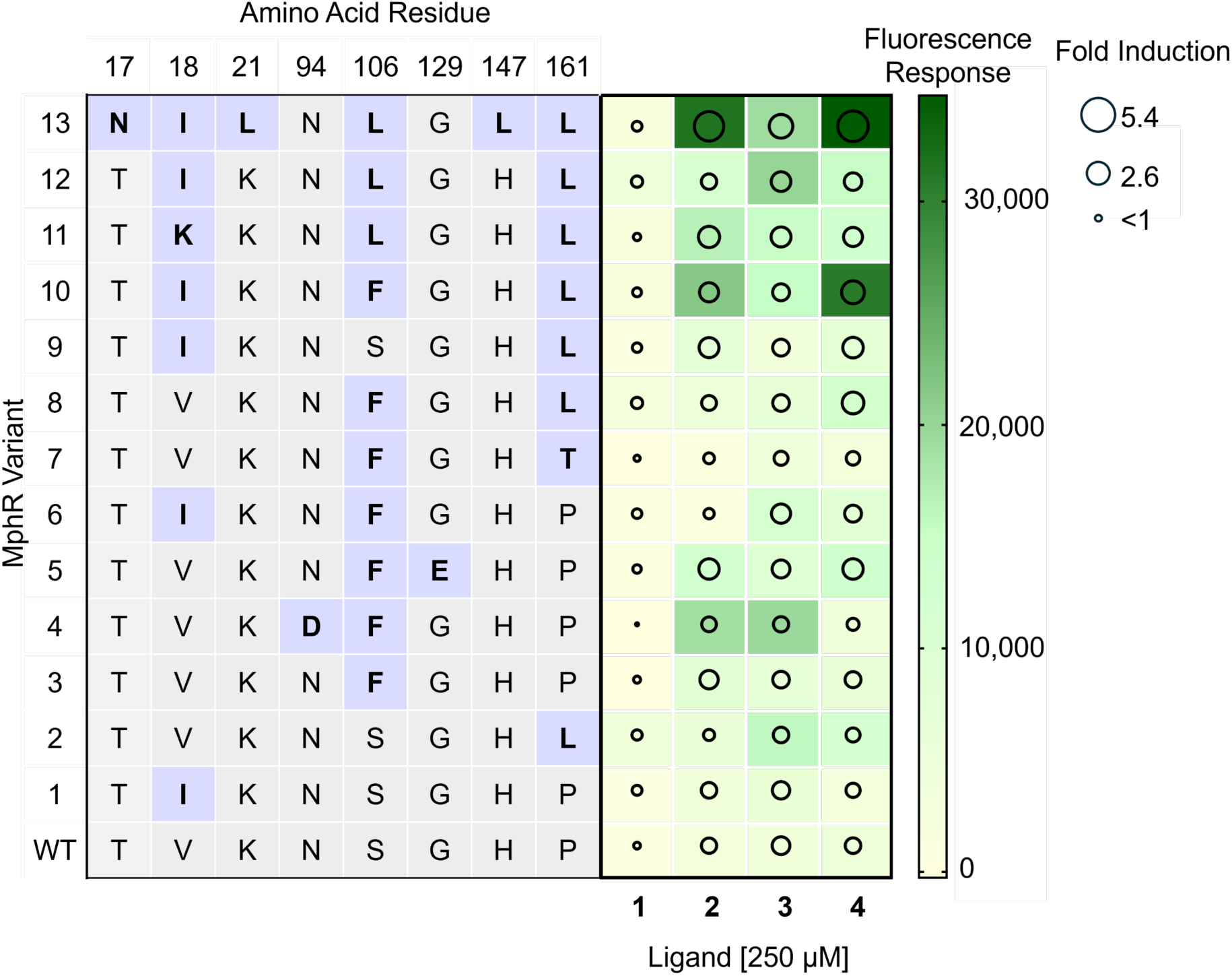
The response of each MphR variant to 250 µM ligand. Amino acid residues bolded with a purple square indicate a new mutation. The color scale indicates relative fluorescence units (RFU) for each pairing. Bubble size indicates the fold induction value.

The MphR mutants used in the initial campaign were selected based on two criteria: effector-structure similarity and observed promiscuity. Previously described wild-type (WT) MphR and its engineered variants detect a variety of 12-, 14-, 15-, and 16-membered macrolides and macrolactones, ranging from the fully decorated 14-membered erythromycin A (ErA) to its aglycone, erythronolide B (EB) [27–29].

The broad range of effectors, along with the ability of more recent MphR mutants to detect the 14-membered aglycones, supports the hypothesis that MphR may be leveraged as a sesquiterpene biosensor.

### 2.2. Multi-Effector Profiling of MphR Variants

To assess each MphR variant’s ability to detect the sesquiterpene panel, we measured superfolder Green Fluorescent Protein (*sf*GFP) expression from the MphR biosensor gene circuit in the presence of each sesquiterpene at a fixed concentration. We housed the biosensors in the *tolC* knockout strain, *E. coli* JW5503, because deleting TolC improves retention of otherwise diffusible effectors within the cell [29]. We *used* the sfGFP measurements to derive three parameters describing biosensor performance. The ‘normalized fluorescence’ (*sf*GFP/OD_600_) simply normalized the fluorescent output to the culture density. The biosensor response for each ligand was calculated as the difference between induced and background normalized fluorescence (RFU), while fold induction was calculated as the ratio of induced to background normalized fluorescence, providing a measure of induction for each sesquiterpene-biosensor pair. As anticipated, WT MphR showed a poor response to the sesquiterpene panel (**Fig. 2**), consistent with the observation that WT MphR largely depends on effectors that include phosphorylated desosamine, found on ErA and other macrolide effectors [28]. The single residue variants (M1, M2, and M3) showed relatively low fluorescence output, with M2 displaying the highest response (15,800 ± 2,000 RFU with **3**). Yet, with a fold induction value of 1.8 ± 0.05, the response of this MphR variant was too low for further consideration. The double mutants (M4-M9) displayed a similarly poor response to the sesquiterpene panel, with M4 displaying the highest response (19,500 ± 500 RFU with 250 µM **2**), but again with a poor fold induction (1.8 ± 0.03), driven by high background fluorescence (**Table S1**).

The triple-mutant variants containing combinations of mutations at positions 18, 106, and 161 (M10-M12) begin to show both high fold induction and fluorescence, with responses for the oxygenated terpenes ranging from 15,000 to 20,000 RFU and fold inductions nearing 3. However, MphR-13, which contains additional mutations at residues 17, 21, and 147, displayed the highest response and fold-induction responses among the panel for the oxygenated terpenes. For instance, **1** supported a response and fold induction of ∼9,500 RFU and ∼1.2, respectively, whereas **2**, **3**, and **4** produced responses of ∼31,000 RFU, ∼20,000 RFU, and ∼35,000 RFU and fold inductions of ∼5.0, ∼3.5, and ∼5.4, respectively. These data suggest that later-generation MphR mutations improve biosensor performance toward sesquiterpenes, and that additional biosensor engineering could further improve the response.

### 2.3. Dose Response Analysis of Wild-Type and MphR-13 with 1-4

To further characterize MphR-13 responses to each sesquiterpene, we generated WT MphR and MphR-13 fluorescence dose-response curves across a range of effector concentrations. We again normalized the fluorescence response to culture density and subtracted the fluorescence response in the absence of ligand to determine induction at each concentration. The dose response data were then fit to the Hill Equation (**Eqn. 1**, see Methods), and the half-maximal effector concentration (*K*_1/2_) and dynamic range were extracted (**Table S6**).

As expected, the response of the WT MphR was minimal, matching what was previously observed (**Fig. 3**). MphR-13 demonstrated a more sensitive response to the oxygenated terpenes **2**-**4** than the WT biosensor strain (**Fig. 3**). MphR-13 was most sensitive to **2**, with a *K*_1/2_ value of 80 ± 6 µM. In addition, **1** and **4** displayed *K*_1/2_ values of ∼830 and ∼110, respectively (**Table S2**). Although MphR-13 showed the highest sensitivity to **2**, as judged by the *K*_1/2_, **4** provided the highest dynamic range at ∼47,000 RFU, compared to ∼42,000 RFU for **1** and ∼25,000 RFU for **2**. While the response of **3** failed to plateau and therefore did not fit the Hill Equation, it reached a maximum response of ∼30,000 RFU. These differences in biosensor performance suggest that each effector may elicit a different allosteric response.

**Figure 3.**
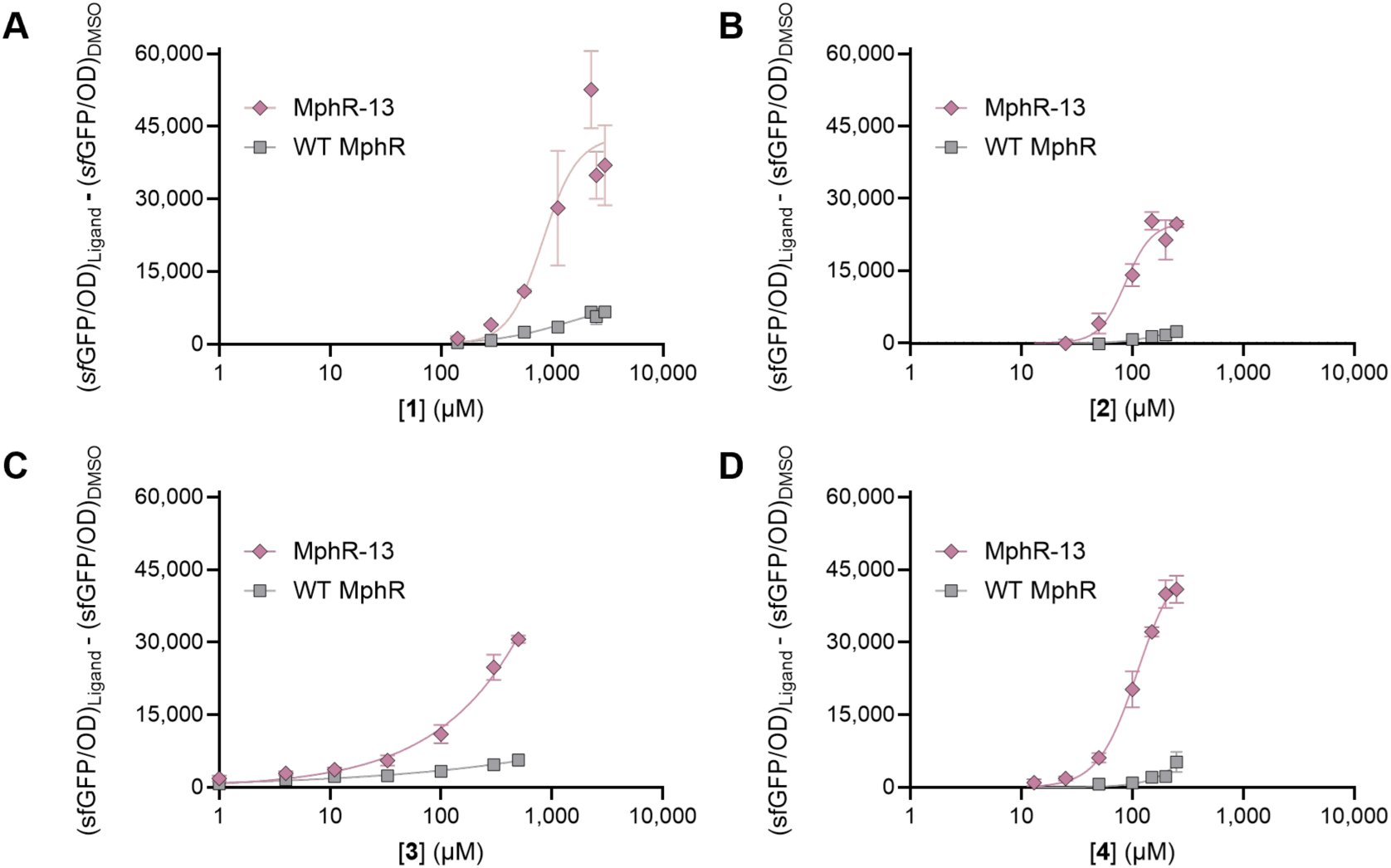
Dose response curves of WT MphR and MphR-13 at various concentrations of **1** (**A**), **2** (**B**), **3** (**C**), and **4** (**D**). Error bars (where visible) represent the standard deviation from the mean, n=3.

Interestingly, **1** elicited only a weak response when initially tested with the MphR panel at 250 µM but induced significant fluorescence with MphR-13 at higher ligand concentrations (>3 mM). At the same **1** concentration, the response of WT MphR was 5.5-fold lower, indicating that the MphR-13 mutations dramatically affect MphR effector promiscuity.

### 2.4. The Role of Individual Mutations in MphR-13 Probed via Site-Directed Mutagenesis

Assays of the initial panel of MphR variants revealed that mutations at Val18, Ser106, and Pro161 were minimally required to support a fluorescence response to sesquiterpenes **1**-**4**, and additional mutants at other positions significantly improved the response. Accordingly, residues 17, 21, and 147 in MphR-13 (Asn, Leu, Leu, respectively) were each targeted individually by site-directed mutagenesis to restore the corresponding wild-type residue (N17T, L21K, and L147H; MphR-13-A, -B, and -C, respectively) (**Table 2**). Each new mutant was then assayed at three concentrations of each sesquiterpene (**1**-**4**) to determine whether the reversal affected the effector response and to identify which mutations in MphR-13 are minimally required for its effector response.

Each reversal mutant maintained a significant response to every ligand tested, but this was accompanied by a reduction in fluorescence response and fold-induction compared with MphR-13 for most ligand-variant combinations (**Fig. 4**, **Table S3**). For instance, reversing the mutations at residues 17 and 21 (MphR-13-A and MphR-13-B) resulted in a ∼1.5x decrease in fluorescence response with **1**-**3**, but not with **4**. Conversely, introducing H147L (MphR-13-C) to reverse the mutation at that position affects only **4**. This suggests that ligand **4** interacts with the ligand binding pocket differently than ligands **1-3**. Notably, reversing mutations at positions 17 and 21 (T17N/K21L; MphR-13-D) significantly decreased fluorescence response to all four ligands. These two residues are in the predicted DNA-binding domain of MphR and may be crucial for releasing MphR from DNA in response to binding of **1**-**4** [35].

**Figure 4.**
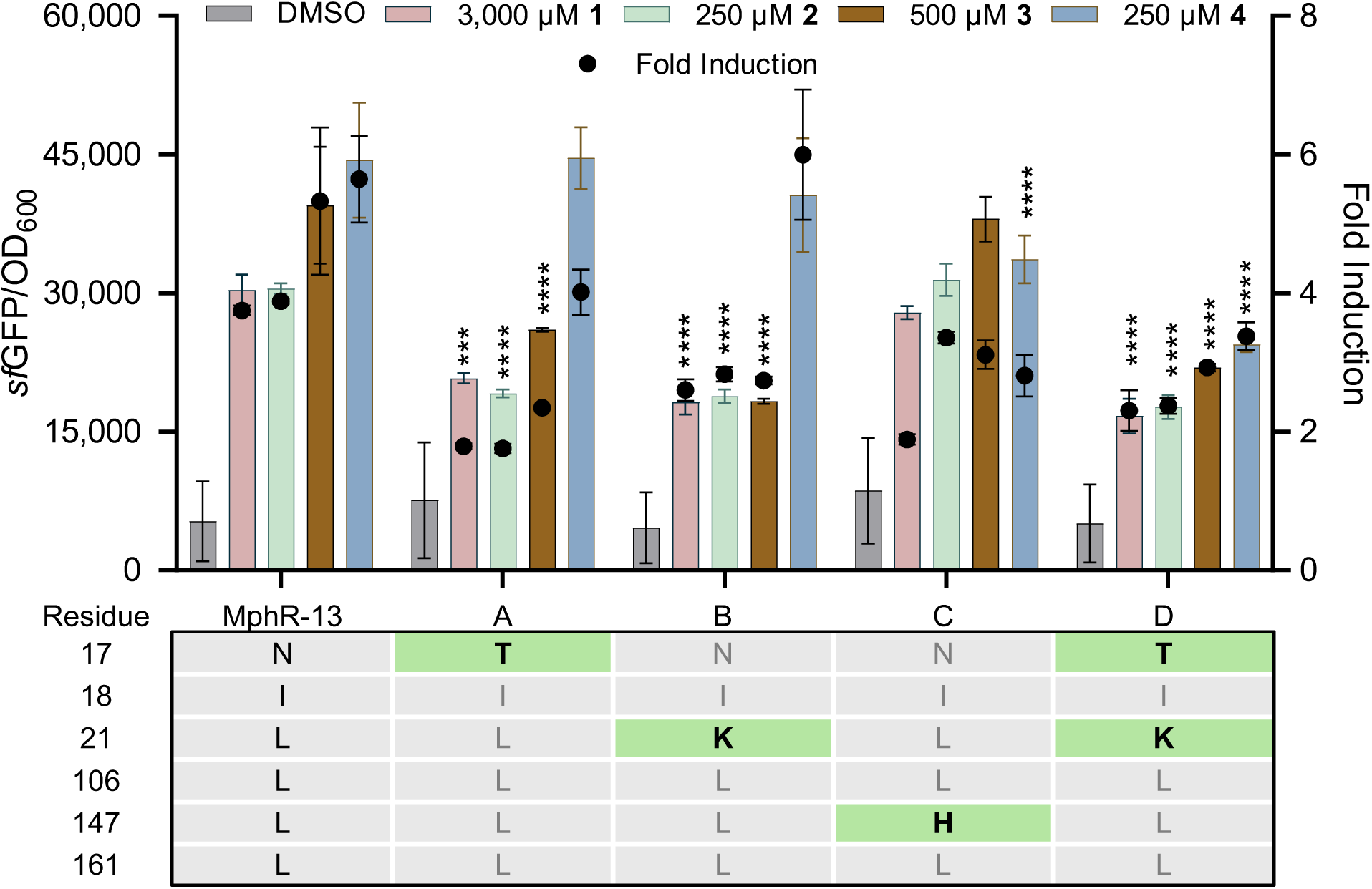
Response of each MphR-13 reversal mutant to the sesquiterpene panel. Each residue highlighted in green in the table indicates that it has been reverted to the WT residue. The normalized fluorescence (bars) is shown on the left Y-axis. The right Y-axis shows the fold induction (normalized fluorescence in the presence of ligand divided by that in the absence of ligand). These values are shown as a single point overlaid on each ligand. Asterisks indicate a significant difference from the NFU of MphR-13 per Tukey’s multiple comparison test (*p*<0.5).

Interestingly, the background fluorescence of each mutant in the absence of **1**-**4** was indistinguishable from that of MphR-13, suggesting that mutations at residues 17, 21, and 147 have limited impact on interactions between the DNA-binding domain and the operator *mphO*; however, further analysis is needed to confirm.

### 2.5. Docking Simulations of MphR-13 with the Sesquiterpene Panel

Each ligand (**1**-**4**) was docked with an AlphaFold-generated model of MphR-13 (**Fig. 5**). Each of the sesquiterpenes was found to dock within the canonical ErA-binding pocket of MphR, with many amino acids previously demonstrated to impact specificity and response residing within 4.5 Å of each ligand, such as Phe99, Arg122, and Asn123 [27,28]. Among the sites mutated in MphR-13, Leu106 is within 4.5 Å of ligands **1** and **3**, whereas Leu147 is within 4.5 Å of **2** and **4**. Ligand **3** shows polar interactions between the oxygen atoms in its 6-and 7-membered rings and the side chain of Arg122, likely forming hydrogen bonds between the oxygen lone pairs and the hydrogen atoms of the side chain amine groups. The predicted binding of **4** involves a hydrogen bond interaction between the **4**-hydroxyl group and the

**Figure 5.**
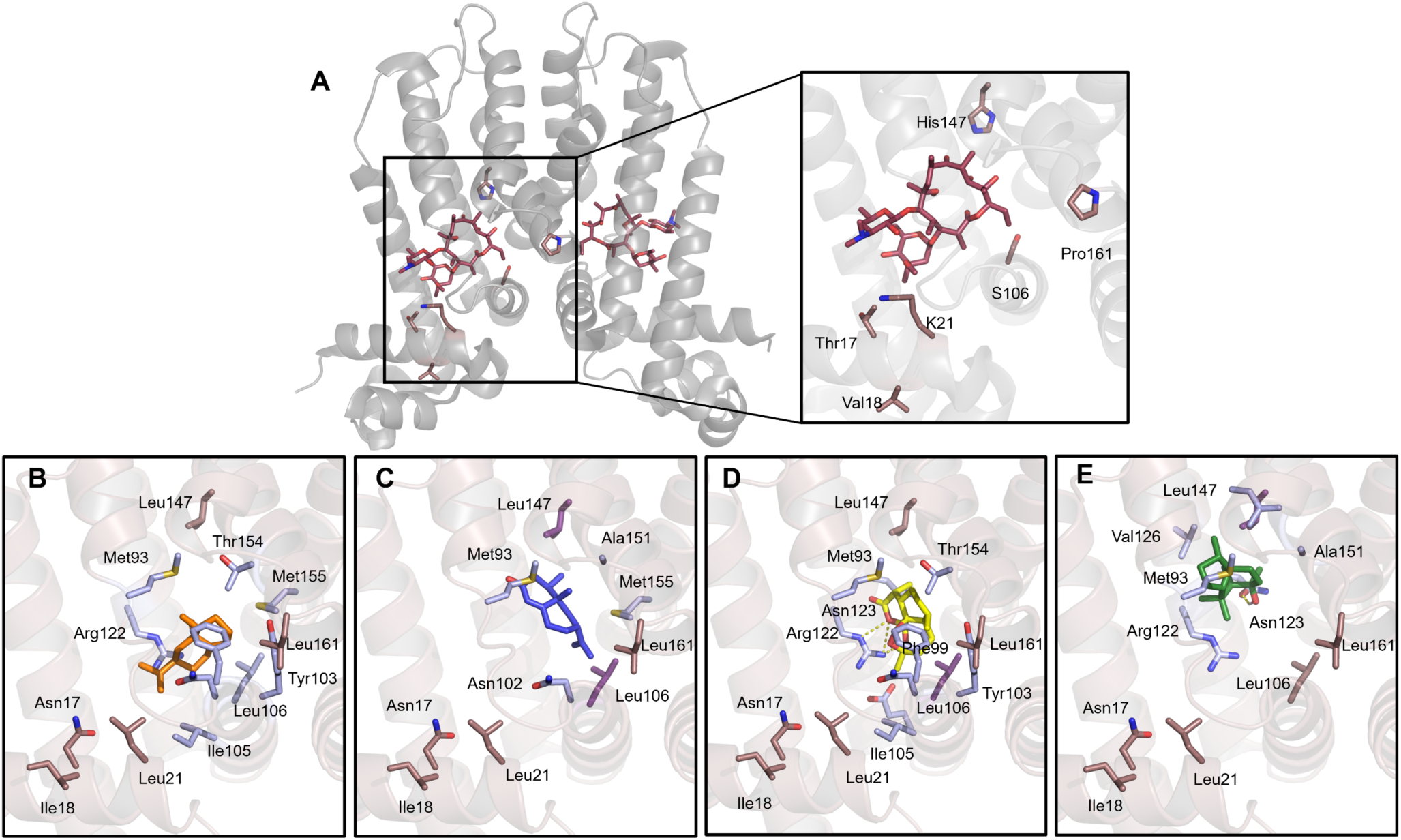
Predicted binding interactions between MphR and 1-4. (**A**) Crystal structure of WT MphR (PDB: 6u18) with ErA. Residues highlighted in light pink represent residues mutated in the MphR-13 mutant. (**D**-**E**) Docking simulations of MphR-13 with **1** (**B**) shown as orange sticks, **2** (**C**) shown as blue sticks, **3** (**D**) shown as yellow sticks, and **4** (**E**) shown as green sticks. Pink sticks represent MphR-13 mutations outside 4.5 Å of the ligand, whereas purple sticks represent MphR-13 mutations within 4.5 Å of the ligand. Blue sticks represent residues that are not mutated within the 4.5 Å range.

Asn123 amide side chain. These results suggest that further investigation of residues around the sesquiterpene ligand panel may affect biosensor performance, such as sensitivity and selectivity.

Computational determination of the ligand binding pockets of WT MphR and MphR-13 reveals that the introduction of the MphR-13 mutations decreases the ligand binding pocket size from an average of 1,880 Å^3^ to 1,640 Å^3^(**Fig. 6A**, **Table S4**). Furthermore, fluorescence output of the MphR biosensors at the highest screened concentration correlated with a decrease in ligand-binding pocket size (**Fig. 6B**). The computed molecular surface of ErA spans 796 Å^2^, whereas that of **1**-**4** is 278, 281, 299, and 286 Å^2^, respectively. This discrepancy, along with the correlation between biosensor pocket size and fluorescence response, suggests that WT MphR cannot respond to sesquiterpenes because its large ligand-binding domain fails to trigger MphR release from DNA. Additionally, replacing many polar residues with nonpolar ones in the engineered MphR variants (e.g., replacing Ser106 with Leu) likely favors binding of smaller, more hydrophobic ligands than ErA.

**Figure 6.**
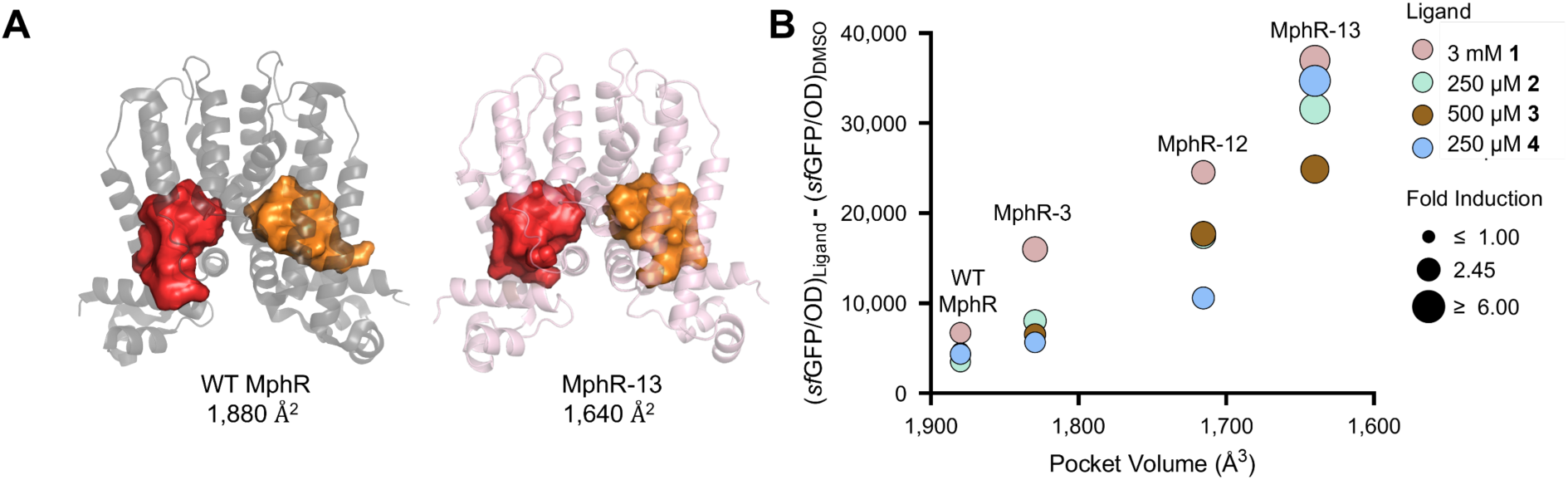
Analysis of the ligand pockets of WT and mutant MphR. (**A**) The pocket volume calculations of WT MphR (*left*) and MphR-13 (*right*). (**B**) A comparison of the pocket volume of MphR mutants and the average fluorescent output in response to each ligand.

### 2.6. Reporter Engineering to Improve the Fold-Induction of MphR-13

The response of transcription factor biosensors has previously been enhanced by replacing the fluorescent reporter and reducing background fluorescence [20]. In the current MphR biosensor construct, the reporter protein is the engineered robust GFP variant, *sf*GFP [28]. However, *sf*GFP has been shown to have high background expression, often deterring from the fold induction values observed in biosensor assays and limiting the dynamic range. In contrast, the αGFP variant has often exhibited high fold induction and low background expression [18]. Accordingly, *sf*GFP in MphR-13 was replaced with the αGFP variant to improve the fold induction of the biosensor strain.

To compare the biosensor performance of MphR-13 αGFP (αMphR-13) and MphR-13 *sf*GFP (*sf*MphR-13), each was tested with ligands **1**-**4** at a fixed concentration, based on their solubility and previously determined ability to derepress MphR. Remarkably, the fold-induction of αMphR-13 with each ligand was 10-25-fold higher than that with *sf*MphR-13 (**Fig. 7**, **Table S2**) due to the dramatic decrease in background fluorescence and the observed increase in response. Subsequently, αGFP is the preferred reporter protein for the sesquiterpene biosensor strains.

**Figure 7.**
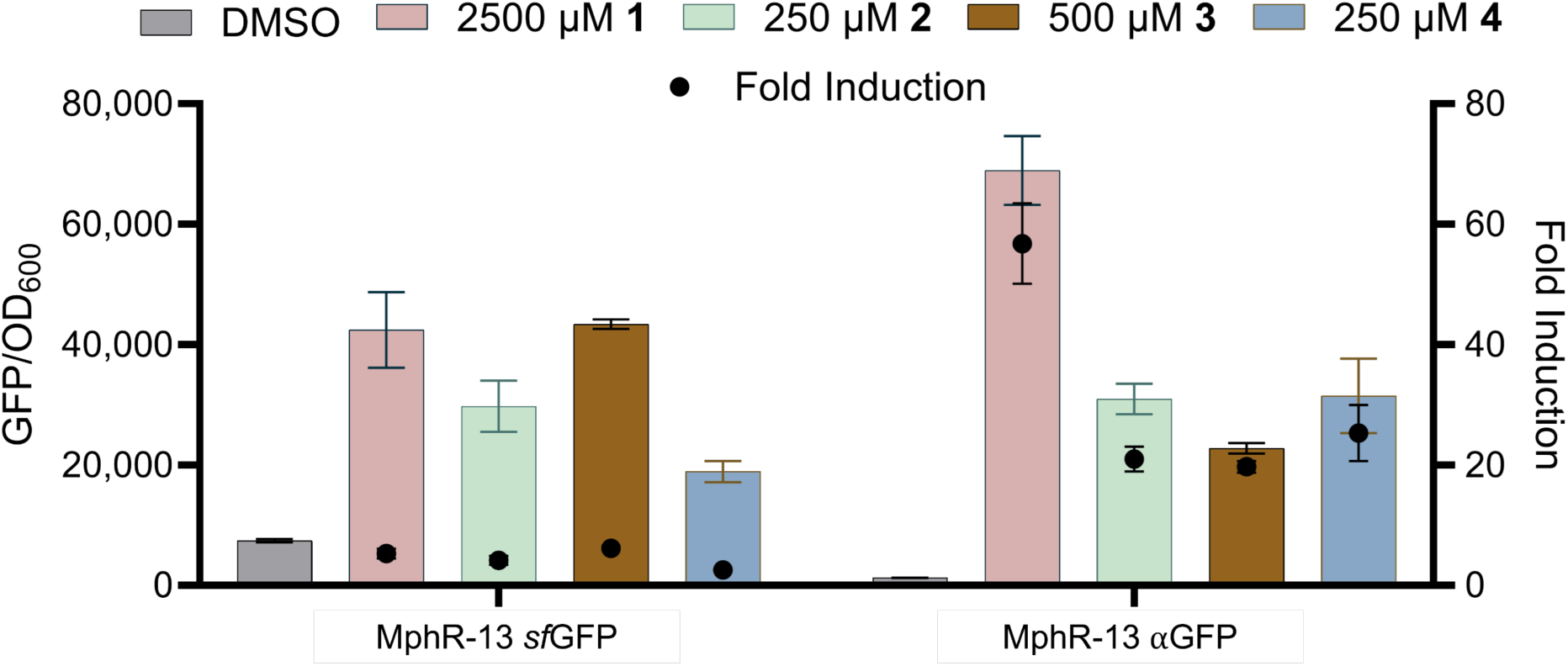
Response of MphR-13 *sf*GFP and MphR-13 αGFP to the sesquiterpene panel. The left Y-axis shows normalized fluorescence (raw fluorescence divided by OD_600_). The right Y-axis shows the fold induction (normalized fluorescence in the presence of ligand divided by that in the absence of ligand). These values are shown as a single point overlaid on each ligand. Error bars represent the standard deviation (n=3).

### 2.7. Probing the Role of the Efflux Pump TolC in Sesquiterpene Detection by MphR

We developed the MphR sesquiterpene assays in the *E. coli* strain JW5503 because deletion of the efflux pump TolC likely increases intracellular ligand concentration and improves the biosensor strain’s sensitivity to aglycone macrolides [29]. To determine whether TolC knock-out improves sesquiterpene detection, we tested and compared the performance of αMphR-13 with **1**-**4** across *E. coli* JW5503, *E. coli* Top10 (negative control), and *E. coli* Top10 Δ*tolC* (**Fig. 8** and **Table S2**). As expected, the JW5503 strain supported the fluorescence response of MphR-13 αGFP with each sesquiterpene tested. Similarly, the Top10 Δ*tolC* strain also supported a robust response, indistinguishable from that of JW5503, except with **3**, which performed ∼2-fold better in the Top10 TolC deletion strain (**Fig. 8** and **Table S2**). In contrast, the Top10 strain failed to elicit a detectable fluorescent response to any ligand across the concentration range tested. These data indicate that *tolC* deletion is absolutely required to detect sesquiterpenes in the *E. coli* biosensor strains.

**Figure 8.**
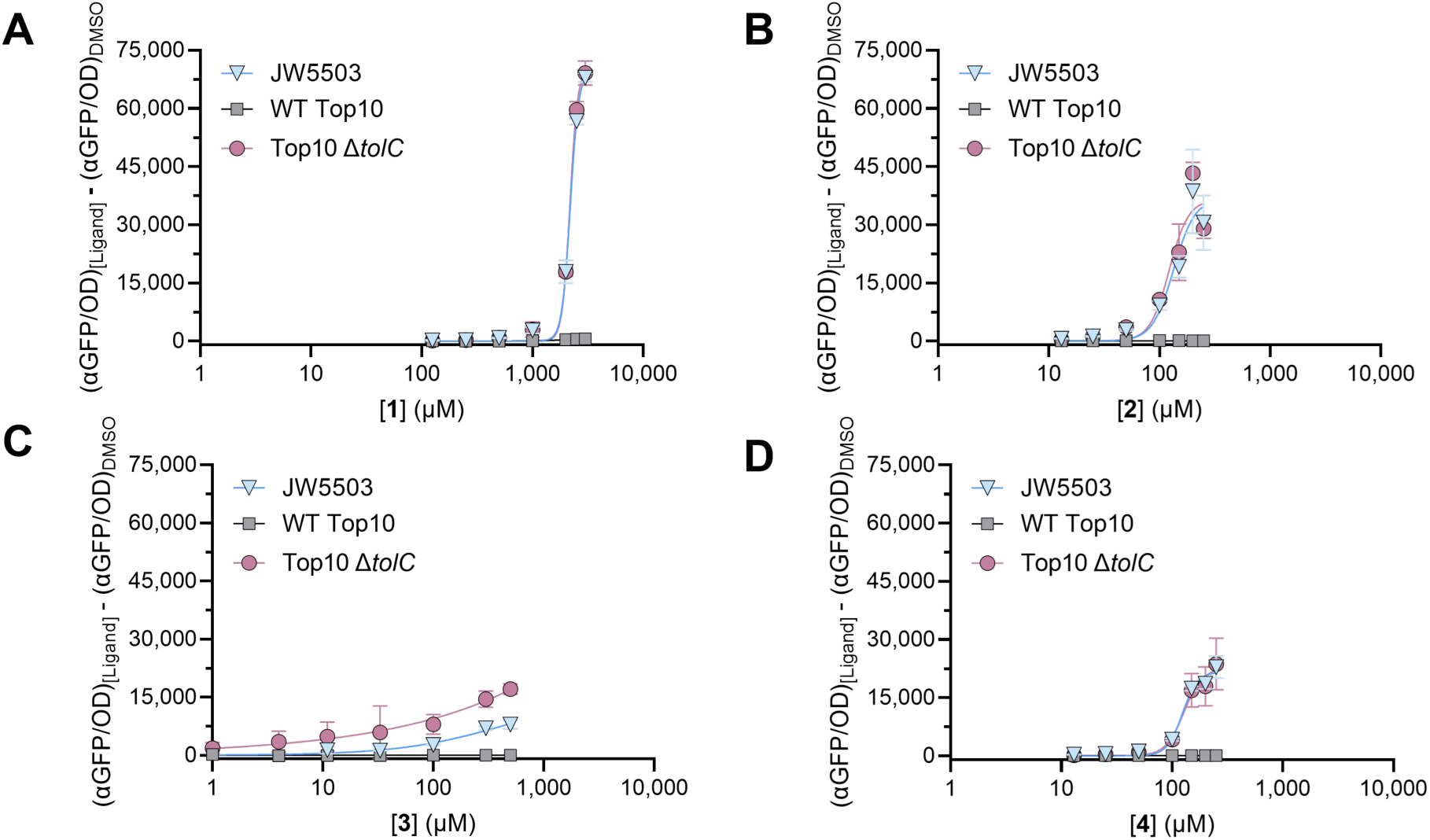
Response of αGFP MphR-13 (αMphR-13) to the sesquiterpene panel in strains in which the gene *tolC* is knocked out. Dose-response curves of JW5503 (triangles), WT Top10 (squares), and Top10 Δ*tolC* (circles) (n=3) with **1** (**A**), **2** (**B**), **3** (**C**), and **4** (**D**). Error bars represent the standard deviation (n=3).

## 3. CONCLUSIONS

Microbial production platforms are highly sought after to provide sustainable, reliable access to valuable terpenes. However, a lack of high-throughput screening methods for *in vivo* terpene production has significantly impeded progress by limiting the number of pathway or enzyme variants that can be tested. In this work, we engineered the transcription factor MphR, which natively responds to large, highly functionalized macrolides, to detect smaller, 15-carbon sesquiterpenes with varying functionalities. We screened a large panel of MphR mutants using four selected sesquiterpenes: valencene (**1**), nootkatone (**2**), artemisinin (**3**), and cedrol (**4**). We used a structure-guided approach to select MphR mutants previously evolved to detect non-phosphorylated macrolides and their aglycones. After screening thirteen MphR variants, we found that six mutations in MphR-13 (T17N/V18I/K21L/S106L/H147L/P161L) transformed the minimal response of the WT transcription factor into an effective sesquiterpene biosensor. Structural studies suggested that these beneficial mutations reshaped the ligand-binding pocket to accommodate smaller, hydrophobic effectors and trigger MphR release from the operator sequence. Yet, the low fold induction of MphR-13 (3-5x) hindered its use as a tool for screening terpene-producing strain libraries. Because the biosensor response is dictated by the entire genetic circuit, we replaced the GFP reporter *sf*GFP with the variant αGFP, improving the signal-to-noise ratio by over 10-fold. Additionally, we showed that knockout of the membrane protein TolC was required for sesquiterpene detection. This suggests that host engineering may be an underutilized tool for improving other genetically encoded terpene biosensors.

MphR has proven to be a highly evolvable and promiscuous transcription factor, providing a platform for developing a sesquiterpene terpene biosensor. The effector specificity of MphR appears to be easily manipulated by directed evolution, suggesting that identifying promiscuous transcription factor biosensors is a promising approach to developing new biosensors [15,18,21,28]. Currently, the ligand promiscuity of αMphR-13 can be leveraged to screen (*1*) total product titers, (*2*) immediate products of terpene cyclases, (*3*) pathways with minimal cyclic intermediates, or (*4*) new sesquiterpene scaffolds. Although αMphR-13 can detect all of **1**-**4**, this mutant is not a true “generalist” sensor, as each sesquiterpene has distinct *K*_1/2_ and fold-induction values. This, coupled with known examples of biosensor effector specificity engineering, suggests that MphR can be further engineered to detect specific terpenes. This would allow the MphR sensor to screen for the end-products of target biosynthetic pathways without unwanted detection of side products or intermediates [27]. Additionally, the development of monoterpene biosensors that distinguish products of P450-catalyzed oxidation of cyclic monoterpenes suggests that the same could be done with sesquiterpenes [22]. Furthermore, the correlation between terpene ligand size and the MphR binding pocket may be leveraged to target larger terpenes (C20+), such as the diterpene paclitaxel.

The directed evolution of MphR provided a rapid approach to developing sesquiterpene biosensors. Building on the success of monoterpene biosensors [21,22], and in conjunction with simplified terpene precursor pathways like the alcohol dependent hemiterpene production pathway [36,37], our MphR variants provide a screening platform for high-throughput synthetic biology of sesquiterpene biosynthetic pathways, potentially overcoming production bottlenecks and providing sustainable access to valuable terpenes.

## 4. MATERIALS AND METHODS

### 4.1. Bacterial Strains, Plasmids, and Materials

Bacterial strains were purchased from Invitrogen unless otherwise stated (**Table S5**). All plasmids were verified by DNA sequencing (**Table S6**). All DNA was purified with kits from New England Biolabs (Ipswich, MA, USA). Synthetic oligonucleotides were purchased from IDT (Coralville, IA, USA). Restriction enzymes were purchased from New England Biolabs. Effector molecules were purchased from Sigma-Aldrich (**1**) and Tokyo Chemical Industry (**2, 3, 4**). Polymerase chain reactions were conducted using Q5 DNA Polymerase from New England Biolabs (NEB). Chemicals were purchased from Sigma-Aldrich (St. Louis, MO, USA, and Thermo Fisher Scientific (Waltham, MA, USA). All commercial enzymes were used according to the manufacturer’s instructions.

### 4.2. Fluorescence Screening of MphR Variants

Unless otherwise specified, the construction of MphR plasmids may be found in prior publications [27–29]. Constructs of the pSENSE plasmid containing each of the *mphR* variants were transformed into *E. coli* JW5503. A single colony of each transformant was picked and used to inoculate a 5 mL starter culture in Luria Broth (LB) containing ampicillin (100 µg/mL), which was incubated overnight at 37 °C with shaking at 250 rpm. The following day, the starter culture was diluted 1:1000 into fresh LB medium containing ampicillin (100 µg/mL) and incubated under the same conditions.

During the growth period, stock solutions of each ligand were diluted in DMSO to prepare solutions at 100x the working concentration (i.e., if a ligand is to be tested at 500 µM, a 50 mM stock solution is prepared). Because of terpene volatility, ligands were prepared no more than 2 h before use.

After 5 h of growth, when the OD_600_ was between 0.6 and 0.7, a multichannel pipette was used to add 495 µL of each culture to a well of a 96-well deep-well microplate. For a zero-ligand control, 5 µL dimethyl sulfoxide (DMSO) was added to row A of the microplate using a multichannel pipette. Then, 5 µL of ligand was added to the remaining wells at increasing concentrations, resulting in a final concentration of 250 µM. A ThermaSeal RTS was used to secure the plate and limit loss of volatile ligand. The microplate was then incubated at 37 °C with shaking at 350 rpm for 16 h. Each construct-ligand combination was screened in triplicate.

The following day, the plates were centrifuged at 1509 *g* for 10 min to harvest the cells. After discarding the supernatant, the pellets were resuspended in 600 µL of sterile phosphate-buffered saline (PBS). The plates were then sealed and vortexed for 5-7 s to resuspend the cells; then 100 µL of the suspension was added to a clear 96-well plate and a black 96-well plate. On a BioTek Hybrid Synergy 4^®^ plate reader, the clear plate was used to measure OD_600,_ and the black plate was used to measure GFP fluorescence (*sf*GFP = Ex 488 nm, Em 510 nm; αGFP = Ex 400 nm, Em 510 nm). The data were analyzed in GraphPad Prism to compare responses across biosensors.

To generate full dose-response curves for select MphR variants, we repeated the process described above across the full concentration range, depending on the ligand (**Table S7**). The data were then plotted using GraphPad Prism^®^ to form a dose-response curve, and the Hill Equation (Equation 1) was used to determine the ligand concentration required to achieve one-half of the maximum RFU (*K*_1/2_)

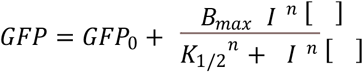

Equation 1: In this equation, GFP is the fluorescent output observed at a given concentration [l]; GFP_0_ refers to the RFU at zero ligand; B_max_ is the maximum fluorescent output observed; *K*_1/2_ is the concentration of ligand that induces ½ the maximum fluorescence; and n is the Hill coefficient.

### 4.3. Site-Directed Mutagenesis of MphR-13

Plasmids containing the MphR-13-reversal mutants were generated via site-directed mutagenesis using Primers P1-P4 (**Table S8**) and Q5 Polymerase. Following agarose gel analysis, the PCR product was repaired using the NEB Kinase-Ligase-DpnI (KLD) enzyme mix. Then, 5 µL of the reaction was transformed into 50 µL of chemically competent *E. coli* Top10 and plated on agar plates containing ampicillin (100 µg/mL). The plasmids were then harvested via the NEB Plasmid Miniprep Kit, and the sequences were confirmed via whole plasmid sequencing. The new constructs were then analyzed via the fluorescence assay described above.

### 4.4. Molecular Modeling of MphR-13 with the Sesquiterpene Panel

A molecular model of the MphR-13 protein was generated using AlphaFold [38]. Each sesquiterpene ligand was then optimized for docking using the MM2 energy minimization function in ChemDraw 3D software, and the resulting structure was saved as a mol2 file. The AlphaFold structure was then opened in PyMOL alongside each of the four sesquiterpenes. The PyMOL plugin DockingPie AutoDock Vina was then used to generate a docking grid box of 19 x 13 x 20, with center coordinates -9.80 x -2.37 x 0.79, to encapsulate the entire MphR-13 protein. Each ligand was then docked using an exhaustiveness of 20 [39,40]. The size of the ligand-binding pockets for the MphR variants was calculated using the PyMOL plugin PyVol [41]. The minimum and maximum radii were set at 1.4 and 5, respectively, whereas the minimum volume was set at 200. The PyMOL “Calculate” action was used to measure the surface area of each ligand in the panel.

### 4.5. Construction of the MphR-13 αGFP Constructs

To replace the gene *sfGFP* with *αGFP* in the MphR-13 construct, Primer P5 was designed to amplify the plasmid region flanking *sfGFP* in MphR-13 (**Table S8**). Additionally, Primer P6 was used to amplify the *αGFP* gene from a plasmid harboring it (**Table S8**). Each PCR product was analyzed on a 1% agarose gel and subsequently purified using a NEB Purification Kit. A Gibson Assembly was then performed according to the manufacturer’s instructions, and the product was immediately transformed into chemically competent Top10 *E. coli*. The DNA sequence of the construct was confirmed by whole-plasmid sequencing.

### 4.6. Construction of ΔtolC Top10 E. coli strain

*E. coli* DH5α containing the plasmid pDK46 was plated on ampicillin-containing agar and incubated overnight at 30 °C. A single colony was then picked and used to inoculate 5 mL of LB broth containing ampicillin, which was then shaken at 225 rpm at 30 °C for 20 h. The plasmid was then purified according to the manufacturer’s instructions. The plasmid was transformed into chemically competent *E. coli* Top10, recovered at 30 °C for 1.5 h, then plated on LB agar with ampicillin and grown overnight. To make electrocompetent *E. coli* Top10 pDK46, a single colony was used to inoculate a 5 mL starter culture in LB broth containing ampicillin. The following day, 100 µL was diluted into 10 mL of fresh media/ampicillin and shaken at 225 rpm and 30 °C until the culture reached an OD_600_ of 0.2, at which point the plasmid was induced with 10 mM L-arabinose. The culture continued to grow under the same conditions until it reached an OD_600_ of 0.6. The culture was then washed twice with 10 mL and 5 mL of 10% glycerol, then resuspended in 2 mL of glycerol and dispersed into 50 µL aliquots. Q5 DNA polymerase and an empty pET28a plasmid were used to amplify the kanamycin resistance gene, *kanR,* using Primer P7 (**Table S8**), which was designed to contain overhangs that complement the DNA flanking the *tolC* gene in the genome.

To perform the deletion of *tolC,* 100 ng of the *kanR* PCR fragment was added to 30 µL of electrocompetent pDK46 T10 *E. coli*. The mixture was then added to a 0.1 mm Gene Pulser Electroporation Cuvette and electroporated at 1.8 kV, 200 Ω, 25 µF using a Gene Pulser Xcell. The cells were immediately suspended in 950 µL of Super Optimal broth with Catabolite repression (S.O.C.) medium and recovered at 225 rpm and 30 °C for 2 h, then at 37 °C for 2 h for pkD46 curing. The transformation was centrifuged at maximum speed for 1 min, and the supernatant was discarded. The cell pellet was resuspended in the remaining volume of SOC by gently pipetting. 50 µL of the transformation was plated on LB agar containing kanamycin (50 ng/µL), and the remaining volume was used to inoculate LB agar containing kanamycin, which was then incubated at 37 °C overnight. Confirmation of the integration of the *kanR* gene was performed via PCR as previously described [42].

## Authorship Contributions

C.L.O. and G.J.W. conceived and designed the experiments. C.L.O. performed the experiments and analyzed the data. A.M.M. and S.C. contributed to experimental design. C.L.O. and G.J.W. wrote the manuscript. All authors read and approved the final manuscript.

## CRediT Authorship Contributions Statement

**Cassidy Oliverio:** Conceptualization, Methodology, Validation, Formal analysis, Investigation, Writing – Original Draft, Visualization; **Alissa Meo:** Conceptualization, Investigation, Resources; **Stefanie Chen:** Investigation, Methodology, Resources; **Gavin Williams:** Conceptualization, Supervision, Project administration, Funding acquisition.

## Acknowledgements

These authors would like to acknowledge Samuel T. Doak, Dongju Yun, and Chris Gupton from the NC State course, BIT 495, “Special Topics in Biotechnology: Directed Evolution”, for the contribution of MphR mutants MphR-5, -7, and -10.

## Appendix A. Supplementary Material

The data supporting this article are included in the supplementary information (SI). Supplementary information: Tables S1-S8 and Fig. S1.

## Data Availability

Raw data files are freely available upon request.

