## Supplemental Information for "Engineering the effector specificity of the macrolide-sensing transcription factor MphR for the detection of sesquiterpenes"

Table of Contents

Supplemental Tables

Table S1: Effector profiling of MphR variants.

Table S2: Hill Equation parameters for WT MphR and MphR-13.

Table S3: Deconvolution of MphR-13 mutations.

Table S4: MphR ligand binding pocket size compared to the biosensor fluorescence output.

Table S5: E. coli strain information.

Table S6: Plasmid information.

Table S7: Ligand dose response ranges

Table S8: Oligonucleotides used in this study.

Figure S1: Surface areas of ErA and ligands 1-4

**SI Table 1**: Performance features of MphR variants

| MphR  Variant | Mutations | Reporter Protein | Uninduced Fluorescence*^a^* | Ligand | RFU*^b^* | Fold Induction***^c^*** |
| --- | --- | --- | --- | --- | --- | --- |
| WT MphR | None | *sf*GFP | 4,500 ± 140 | **1** | 520 ± 20 | 1.03 ± 0.05 |
|  |  |  |  | **2** | 3,390 ± 1,410 | 1.83 ± 0.56 |
|  |  |  |  | **3** | 3,950 ± 210 | 2.23 ± 0.08 |
|  |  |  |  | **4** | 4,770 ± 480 | 1.88 ± 0.09 |
| MphR-1 | V18I | *sf*GFP | 6,300 ± 530 | **1** | 360 ± 410 | 1.23 ± 0.03 |
|  |  |  |  | **2** | 3,890 ± 820 | 1.87 ± 0.06 |
|  |  |  |  | **3** | 5,480 ± 340 | 2.16 ± 0.05 |
|  |  |  |  | **4** | 2,840 ± 580 | 1.68 ± 0.04 |
| MphR-2 | P161L | *sf*GFP | 19,600 ± 1,040 | **1** | 4,140 ± 820 | 1.25 ± 0.02 |
|  |  |  |  | **2** | 5,680 ± 770 | 1.33 ± 0.12 |
|  |  |  |  | **3** | 15,850 ± 1,350 | 1.87 ± 0.06 |
|  |  |  |  | **4** | 11,850 ± 620 | 1.66 ± 0.12 |
| MphR-3 | S106F | *sf*GFP | 6,000 ± 20 | **1** | 140 ± 40 | 1.04 ± 0.01 |
|  |  |  |  | **2** | 7,960 ± 340 | 2.34 ± 0.09 |
|  |  |  |  | **3** | 6,490 ± 220 | 2.04 ± 0.14 |
|  |  |  |  | **4** | 5,680 ± 590 | 1.93 ± 0.11 |
| MphR-4 | N94D/S106F | *sf*GFP | 21,500 ± 1,060 | **1** | -270 ± 550 | 0.94 ± 0.04 |
|  |  |  |  | **2** | 18,760 ± 1,490 | 1.75 ± 0.01 |
|  |  |  |  | **3** | 19,560 ± 870 | 1.83 ± 0.03 |
|  |  |  |  | **4** | 4,170 ± 5,970 | 1.43 ± 0.05 |
| MphR-5 | S106F/E129E | *sf*GFP | 7,500 ± 120 | **1** | 780 ± 80 | 1.12 ± 0.02 |
|  |  |  |  | **2** | 12,860 ± 790 | 2.72 ± 0.22 |
|  |  |  |  | **3** | 9,180 ± 200 | 2.25 ± 0.02 |
|  |  |  |  | **4** | 13,340 ± 2,180 | 2.73 ± 0.28 |
| MphR-6 | S106F/P161T | *sf*GFP | 9,400 ± 320 | **1** | 860 ± 510 | 1.2 ± 0.03 |
|  |  |  |  | **2** | 1,130 ± 720 | 1.26 ± 0.07 |
|  |  |  |  | **3** | 11,790 ± 900 | 2.48 ± 0.03 |
|  |  |  |  | **4** | 8,260 ± 2,750 | 2.08 ± 0.4 |
| MphR-7 | S106F/P161L | *sf*GFP | 4,900 ± 190 | **1** | -240 ± 440 | 1 ± 0.07 |
|  |  |  |  | **2** | 740 ± 220 | 1.3 ± 0.2 |
|  |  |  |  | **3** | 4,240 ± 230 | 1.66 ± 0.08 |
|  |  |  |  | **4** | 3,150 ± 1,170 | 1.6 ± 0.25 |
| MphR-8 | V18I/SP161L | *sf*GFP | 5,600 ± 340 | **1** | 2,330 ± 260 | 1.35 ± 0.03 |
|  |  |  |  | **2** | 4,760 ± 610 | 1.77 ± 0.16 |
|  |  |  |  | **3** | 6,460 ± 1,390 | 2.04 ± 0.21 |
|  |  |  |  | **4** | 12,280 ± 5,530 | 3.02 ± 0.82 |
| MphR-9 | V18I/S106F | *sf*GFP | 5,300 ± 200 | **1** | 330 ± 2,960 | 1.19 ± 0.16 |
|  |  |  |  | **2** | 7,230 ± 530 | 2.56 ± 0.24 |
|  |  |  |  | **3** | 4,020 ± 1,240 | 2.06 ± 0.12 |
|  |  |  |  | **4** | 7,010 ± 230 | 2.76 ± 0.26 |
| MphR-10 | V18I/S106F/  P161L | *sf*GFP | 13,800 ± 170 | **1** | 1,310 ± 370 | 1.15 ± 0 |
|  |  |  |  | **2** | 21,530 ± 670 | 2.47 ± 0.04 |
|  |  |  |  | **3** | 15,330 ± 760 | 2.08 ± 0.01 |
|  |  |  |  | **4** | 30,720 ± 5,390 | 3.27 ± 0.46 |
| MphR-11 | V18K/S106F/  P161L | *sf*GFP | 9,500 ± 160 | **1** | 1,250 ± 400 | 1.06 ± 0.08 |
|  |  |  |  | **2** | 16,740 ± 880 | 2.9 ± 0.11 |
|  |  |  |  | **3** | 15,170 ± 1,190 | 2.63 ± 0.12 |
|  |  |  |  | **4** | 13,550 ± 1,510 | 2.45 ± 0.21 |
| MphR-12 | V18I/F106L/  P161L | *sf*GFP | 11,300 ± 1,310 | **1** | 5,100 ± 990 | 1.35 ± 0.03 |
|  |  |  |  | **2** | 10,760 ± 1,340 | 1.97 ± 0.2 |
|  |  |  |  | **3** | 19,920 ± 1,490 | 2.57 ± 0.06 |
|  |  |  |  | **4** | 15,240 ± 2,280 | 2.18 ± 0.24 |
| MphR-13 | T17N/V18I/  K21L/S106L/H147L/P161L | *sf*GFP | 7,800 ± 230 | **1** | 1,670 ± 360 | 1.22 ± 0.15 |
|  |  |  |  | **2** | 31,550 ± 2,770 | 5.05 ± 0.29 |
|  |  |  |  | **3** | 19,090 ± 100 | 3.45 ± 0.07 |
|  |  |  |  | **4** | 34,730 ± 11,500 | 5.4 ± 1.41 |

*^a^*[*sf*GFP/OD_600_]_DMSO_

*^b^*[*sf*GFP/OD_600_]_Ligand_ - [*sf*GFP/OD_600_]_DMSO_ (at 250 µM ligand)

^c^ [sfGFP/OD_600_]_Ligand_ / [sfGFP/OD_600_]_DMSO_

**Table S2:** Hill Equation performance features of WT MphR and MphR-13

| Strain | MphR Variant | Mutations | Reporter Protein | Uninduced *^a^* | Ligand | B_max_ *^b^* | n *^c^* | *K*_1/2_ *^d^* | Fold Induction *^e^* |
| --- | --- | --- | --- | --- | --- | --- | --- | --- | --- |
| JW5503 |  | WT MphR | *sf*GFP | 4,500 ± 140 | 1 | 9,300 ± 4,200 | 1.29 ± 0.56 | 1,440 ± 1190 | 2.47 ± 0.62 |
|  |  |  |  |  | 2 | 2,900 ± 1,600 | 2.93 ± 2.21 | 160 ± 80 | 1.38 ± 0.14 |
|  |  |  |  |  | 3 | -*^f^* |  |  | 2.11 ± 0.44 |
|  |  |  |  |  | 4 |  |  |  | 1.94 ± 0.31 |
| JW5503 | MphR-13 | T17N/ V18I/ K21L/ S106L/ H147L/ P161L | sfGFP |  | 1 | 42,800 ± 4,300 | 2.78 ± 1.1 | 830 ± 140 | 4.97 ± 0.9 |
|  |  |  |  |  | 2 | 25,000 ± 1,500 | 3.65 ± 0.89 | 90 ± 10 | 3.37 ± 0.71 |
|  |  |  |  |  | 3 | 15,200 ± 1,200 | 4.03 ± 1.4 | 80 ± 10 | 6.18 ± 0.24 |
|  |  |  |  |  | 4 | 47,800 ± 2,900 | 2.44 ± 0.28 | 110 ± 10 | 5.54 ± 0.49 |
| JW5503 | MphR-13 | T17N/ V18I/ K21L/ S106L/ H147L/ P161L | αGFP | 1,300 ± 10 | 1 | 76,000 ± 3,600 | 27 ± 13 | 2320 ± 90 | 56.78 ± 5.44 |
|  |  |  |  |  | 2 | 36,800 ± 5,200 | 4.52 ± 1.72 | 130 ± 10 | 21 ± 1.67 |
|  |  |  |  |  | 3 | 40,700 ± 7,700 | 3.13 ± 0.64 | 180 ± 30 | 19.7 ± 0.78 |
|  |  |  |  |  | 4 | 21,600 ± 800 | 6.32 ± 0.93 | 120 ± 0 | 20.12 ± 0.82 |
| Top10 | MphR-13 | T17N/ V18I/ K21L/ S106L/ H147L/ P161L | αGFP | 700 ± 60 | 1 | 700 ± 100 | 3.44 ± 0.96 | 2,060 ± 290 | 1.72 ± 0.17 |
|  |  |  |  |  | 2 |  |  |  | 1.09 ± 0.08 |
|  |  |  |  |  | 3 |  |  |  | 1.08 ± 0.36 |
|  |  |  |  |  | 4 |  |  |  | 1.03 ± 0.02 |
| Δ*tolC* Top10 | MphR-13 | T17N/ V18I/ K21L/ S106L/ H147L/ P161L | αGFP | 1,700 ± 140 | 1 | 70,500 ± 1,400 | 12.49 ± 0.84 | 2,180 ± 20 | 40.52 ± 2.48 |
|  |  |  |  |  | 2 | 36,600 ± 3,900 | 4.98 ± 1.82 | 120 ± 10 | 19.91 ± 1.98 |
|  |  |  |  |  | 3 | 3,784,100 ± | 0.37 ± 0.25 |  | 10.09 ± 2.15 |
|  |  |  |  |  | 4 | 22,600 ± 2,300 | 5.24 ± 1.74 |  | 18.02 ± 4.03 |

*^a^* The Raw Fluorescence/OD_600_ of the biosensor in the absence of ligand

*^b^* When the uninduced fluorescence subtracted, GFP_Max_ – GFP_min_

*^c^* Hill coefficient that measures the cooperativity of ligand binding. N>1 indicates a positive binding

*^d^* The concentration of ligand at 1/2 the maximum fluorescence output

*^e^* The normalized fluorescence output of the sensor in response to the highest screened concentration of ligand divided by the uninduced fluorescence

^f^ Not Calculated (N.C.) indicates that while the biosensor had a response, the data was unable to reach a maximum fluorescence and therefore was unable to fit to the Hill equation properly.

**SI Table 3**: Performance features of MphR variants related to deconvolution of MphR-13

|  | MphR Variant | Reporter Protein | Uninduced Fluorescence *^a^* | Ligand | Maximum RFU *^b^* | Fold Induction *^c^* |
| --- | --- | --- | --- | --- | --- | --- |
| MphR-13 | T17N/V18I/  K21L/S106L/ H147L/P161L | *sf*GFP | 7,800 ± 300 | **1** | 30,300 ± 1,200 | 3.75 ± 0.05 |
|  |  |  |  | **2** | 30,500 ± 500 | 3.88 ± 0.03 |
|  |  |  |  | **3** | 39,500 ± 5,200 | 5.33 ± 0.87 |
|  |  |  |  | **4** | 44,400 ± 4,400 | 5.64 ± 0.44 |
| MphR-13a | V18I/K21L/ S106L/H147L/ P161L | *sf*GFP | 11,200 ± 300 | **1** | 20,800 ± 500 | 1.79 ± 0.03 |
|  |  |  |  | **2** | 19,100 ± 300 | 1.76 ± 0.06 |
|  |  |  |  | **3** | 26,000 ± 100 | 2.34 ± 0.01 |
|  |  |  |  | **4** | 44,600 ± 2,700 | 4.01 ± 0.27 |
| MphR-13b | T17N/V18I/ S106L/H147L/ P161L | *sf*GFP | 6,800 ± 200 | **1** | 18,200 ± 1,100 | 2.6 ± 0.13 |
|  |  |  |  | **2** | 18,800 ± 600 | 2.83 ± 0.08 |
|  |  |  |  | **3** | 18,300 ± 200 | 2.74 ± 0.04 |
|  |  |  |  | **4** | 40,600 ± 5,000 | 6 ± 0.77 |
| MphR-13c | T17N/V18I/ K21L/S106L/ P161L | *sf*GFP | 11,900 ± 2,000 | **1** | 27,900 ± 600 | 1.89 ± 0.06 |
|  |  |  |  | **2** | 31,400 ± 1,400 | 3.36 ± 0.07 |
|  |  |  |  | **3** | 38,000 ± 2,000 | 3.11 ± 0.17 |
|  |  |  |  | **4** | 33,600 ± 2,100 | 2.81 ± 0.24 |
| MphR-13d | V18I/S106L/ H147L/P161L | *sf*GFP | 7,500 ± 200 | **1** | 16,700 ± 1,500 | 2.3 ± 0.24 |
|  |  |  |  | **2** | 17,600 ± 1,000 | 2.37 ± 0.09 |
|  |  |  |  | **3** | 21,900 ± 100 | 2.93 ± 0.04 |
|  |  |  |  | **4** | 24,400 ± 700 | 3.38 ± 0.16 |

^a^[sfGFP/OD_600_]_DMSO_

*^b^*[*sf*GFP/OD_600_]_Ligand_^-^[*sf*GFP/OD_600_]_DMSO_

^c^[sfGFP/OD_600_]_Ligand_/[sfGFP/OD_600_]_DMSO_

**Table S4**: Performance features of MphR variants related to ligand binding pocket size

| MphR Variant | Mutations | Average Pocket Size (Å) | Ligand | Maximum RFU *^a^* | Fold Induction *^b^* |
| --- | --- | --- | --- | --- | --- |
| WT MphR | None | 1,880 | 3 mM **1** | 6,700 ± 260 | 2.47 ± 0.62 |
|  |  |  | 250 µM **2** | 3,500 ± 1,970 | 1.83 ± 0.56 |
|  |  |  | 500 µM **3** | 4,700 ± 200 | 2.23 ± 0.08 |
|  |  |  | 250 µM **4** | 4,300 ± 390 | 1.88 ± 0.09 |
| MphR-3 | S106F | 1,830 | 3 mM **1** | 27,900 ± 4,720 | 3.03 ± 0.17 |
|  |  |  | 250 µM **2** | 8,000 ± 430 | 2.34 ± 0.09 |
|  |  |  | 500 µM **3** | 6,400 ± 510 | 2.04 ± 0.14 |
|  |  |  | 250 µM **4** | 5,600 ± 620 | 1.93 ± 0.11 |
| MphR-12 | V18I/S106L/ P161L | 1,715 | 3 mM **1** | 37,000 ± 6,710 | 4.97 ± 0.9 |
|  |  |  | 250 µM **2** | 11,200 ± 1,000 | 2.23 ± 0.08 |
|  |  |  | 500 µM **3** | 17,700 ± 530 | 3.09 ± 0.15 |
|  |  |  | 250 µM **4** | 10,600 ± 760 | 2.23 ± 0.08 |
| MphR-13 | T17N/V18I/ K21L/S106L H147L/P161L | 1,640 | 3 mM **1** | 17,600 ± 1,240 | 2.82 ± 0.32 |
|  |  |  | 250 µM **2** | 31,600 ± 2,590 | 5.05 ± 0.29 |
|  |  |  | 500 µM **3** | 19,100 ± 80 | 3.45 ± 0.07 |
|  |  |  | 250 µM **4** | 34,700 ± 11,310 | 5.4 ± 1.41 |

*^a^* [*sf*GFP/OD_600_]_Ligand_^-^[*sf*GFP/OD_600_]_DMSO_

^b^[sfGFP/OD_600_]_Ligand_/[sfGFP/OD_600_]_DMSO_

**Table S5**: Genotypes of E. coli strains used in this study

| **Name** | **Genotype** | **Reference** |
| --- | --- | --- |
| Top10 | F^–^*mcr*A Δ(*mrr*-*hsd*RMS-*mcr*BC) φ80*lac*ZΔM15 Δ*lac*X74 *rec*A1 *ara*D139 Δ(*ara-leu*)7697 *gal*U *gal*K λ–*rps*L(Str^R^) *end*A1 *nup*G | Invitrogen |
| JW5503 | F- Δ(araD-araB)567 ΔlacZ4787(::rrnB-3) λΔtolC732::kan rph-1 Δ(rhaD-rhaB)568 hsdR514 | ^53^ |
| Top10 Δ*tolC* | F^–^*mcr*A Δ(*mrr*-*hsd*RMS-*mcr*BC) φ80*lac*ZΔM15 Δ*lac*X74 *rec*A1 *ara*D139 Δ(*ara-leu*)7697 *gal*U *gal*K λ–*rps*L(Str^R^) *end*A1 *nup*G λΔtolC732::kan | This work |

Table S6: Plasmid maps

| Plasmid Name and Color-Coding Scheme | Sequence |
| --- | --- |
| pSENSE-MphR-*sf*GFP  *mphR* SEQUENCE  mphO operator sequence  *lac promoter*  *lacIQ promoter*  *sf*GFP SEQUENCE  Origin of Replication  Ampicillin resistance | gtttgcaagcagcagattacgcgcagaaaaaaaggatctcaagaagatcctttgatcttttctacggggtctgacgctcagtggaacgaaaactcacgttaagggattttggtcatgagattatcaaaaaggatcttcacctagatccttttaaattaaaaatgaagttttaaatcaatctaaagtatccatggatatgagtaaacttggtctgacagttaccaatgcttaatcagtgaggcacctatctcagcgatctgtctatttcgttcatccatagttgcctgactccccgtcgtgtagataactacgatacgggagggcttaccatctggccccagtgctgcaatgataccgcgagacccacgctcaccggctccagatttatcagcaataaaccagccagccggaagggccgagcgcagaagtggtcctgcaactttatccgcctccatccagtctattaattgttgccgggaagctagagtaagtagttcgccagttaatagtttgcgcaacgttgttgccattgctacaggcatcgtggtgtcacgctcgtcgtttggtatggcttcattcagctccggttcccaacgatcaaggcgagttacatgatcccccatgttgtgcaaaaaagcggttagctccttcggtcctccgatcgttgtcagaagtaagttggccgcagtgttatcactcatggttatggcagcactgcataattctcttactgtcatgccatccgtaagatgcttttctgtgactggtgagtactcaaccaagtcattctgagaatagtgtatgcggcgaccgagttgctcttgcccggcgtcaacacgggataataccgcgccacatagcagaactttaaaagtgctcatcattggaaaacgttcttcggggcgaaaactctcaaggatcttaccgctgttgagatccagttcgatgtaacccactcgtgcacccaactgatcttcagcatcttttactttcaccagcgtttctgggtgagcaaaaacaggaaggcaaaatgccgcaaaaaagggaataagggcgacacggaaatgttgaatactcatactcttcctttttcaatattattgaagcatttatcagggttattgtctcatgagcggatacatatttgaatgtatttagaaaaataaacaaaaagagcatgcgtttgtagaaacgcaaaaaggccatccgtcaggatggccttctgcttaatttgatgcctggcagtttatggcgggcgtcctgcccgccaccctccgggccgttgcttcgcaacgttcaaatccgctcccggcggatttgtcctactcaggagagcgttcaccgacaaacaacagataaaacgaaaggcccagtctttcgactgagcctttcgttttatttgatgcctggcagttccctactctcgcatggggagaccccacactaccatcggcgctacggcgtttcacttctgagttcggcatggggtcaggtgggaccaccgcgctactgccgccaggcaaattctgttttatcagaccgcttctgcgttctgatttaatctgtatcaggctgaaaatcttctctcatccgccaaaacagccaagctggagaccgTTTAAACGGGCCCAAGCTTTTTGTAGAGCTCATCCATGCCATGTGTAATCCCAGCAGCAGTTACAAACTCAAGAAGGACCATGTGGTCACGCTTTTCGTTGGGATCTTTCGAAAGGACAGATTGTGTCGACAGGTAATGGTTGTCTGGTAAAAGGACAGGGCCATCGCCAATTGGAGTATTTTGTTGATAATGGTCTGCTAGTTGAACGGAACCATCTTCAACGTTGTGGCGAATTTTGAAGTTAGCTTTGATTCCATTCTTTTGTTTGTCTGCCGTGATGTATACATTGTGTGAGTTAAAGTTGTACTCGAGTTTGTGTCCAAGAATGTTTCCATCTTCTTTAAAATCAATACCCTTTAACTCGATACGATTAACAAGGGTATCACCTTCAAACTTGACTTCAGCACGCGTCTTGTAGGTCCCGTCATCTTTGAAAGATATAGTGCGTTCCTGTACATAACCTTCGGGCATGGCACTCTTGAAAAAGTCATGCCGTTTCATGTGATCCGGATAACGGGAAAAGCATTGAACACCATAGGTCAGAGTAGTGACAAGTGTTGGCCACGGAACAGGTAGTTTTCCAGTAGTGCAAATAAATTTAAGGGTGAGTTTTCCGTTTGTAGCATCACCTTCACCCTCTCCACGGACAGAAAATTTGTGCCCATTAACATCACCATCTAATTCAACAAGAATTGGGACAACTCCAGTGAAAAGTTCTTCTCCTTTGCTCATActagtttcctcctgttagcccacctaaatgtaacagtcacgtcggttatattcaatcccaca*caacatacgagccggaagcataaagtgtaaa*gcctggggtgcctaatgagtgagctaactcacattaattgcgttgcgctctag*atggtgcaaaacctttcgcggtatggcatgatagcgccc*cccaacccatacagaaggtgaacactgATGCCCCGCCCCAAGCTCAAGTCCGATGACGAGGTACTCGAGGCCGCCACCGTAGTGCTGAAGCGTTGCGGTCCCATAGAGTTCACGCTCAGCGGAGTAGCAAAGGAGGTGGGGCTCTCCCGCGCAGCGTTAATCCAGCGCTTCACCAACCGCGATACGCTGCTGGTGAGGATGATGGAGCGCGGCGTCGAGCAGGTGCGGCATTACCTGAATGCGATACCGATAGGCGCAGGGCCGCAAGGGCTCTGGGAATTTTTGCAGGTGCTCGTTCGGAGCATGAACACTCGCAACGACTTCTCGGTGAACTATCTCATCTCCTGGTACGAGCTCCAGGTGCCGGAGCTACGCACGCTTGCGATCCAGCGGAACCGCGCGGTGGTGGAGGGGATCCGCAAGCGACTGCCCCCAGGTGCTCCTGCGGCAGCTGAGTTGCTCCTGCACTCGGTCATCGCTGGCGCGACGATGCAGTGGGCCGTCGATCCGGATGGTGAGCTAGCTGATCATGTGCTGGCTCAGATCGCTGCCATCCTGTGTTTAATGTTTCCCGAACACGACGATTTCCAACTCCTCCAGGCACATGCGTAAcatagtatccaaaataaggctaagaattcatgatgagcagttcggacgagcagccgcgcccgcgtcgccgcaaccaggatcggcagcaccccaaccagaaccggccggtgctgggccgtaccgagcgggaccgcaaccggcgccagttcgggcagaacttcctccgcgaccgcaagaccatcgcgcgcatcgccgagacagccgagctgcggcccgatctgccggtgctggaagccggccccggcgaagggctgctcaccagggaactcgccgaccgcgcgcgtcaggtgacgtcgtacgagatcgacccccggctggcgaagtcgttgcgggagaagctttccggccacccgaacatcgaagtcgtcaacgccgacttcctcaccgccgaaccgccgcccgagccgttcgccttcgtcggcgcgatcccctacggcatcacctcggcgatcgtggactggtgcctggaggcgccgacgatcgagacggcgacgatggtcacgcagctggagttcgcccggaagcggaccggcgattacggccgctggagccgcctcacggtgatgacctggccgctgttcgagtgggagttcgtcgagaaggtcgaccgccggctgttcaagccggtgcccaaggtcgactcggcgatcatgcggctgcgcaggcgcgccgaaccgctgctggaaggcgcggcgctcgaacgctacgagtcgatggtcgagctgtgcttcaccggcgtcgacggcaacatccaggcgtcgcttctgcgcaagtacccgaggcgccgcgtcgaggcggcgctcgaccacgcgggggtcgggggcggcgccgtggtcgcctacgtccggccggagcagtggctccggctgttcgagcggctggatcagaagaacgaaccgaggggtgggcagccccagcggggcaggcgaaccggcggacgggaccacggggaccggcgaaccggcgggcaggatcgcggcgatcggcgaaccggcggccgcgaccacagggaccggcaagccagcggccacggcgatcgtcgcagcagcggacgcaatcgcgacgacggacgaaccggcgagcgcgagcgtggggaccaaggcgggcggcgggggccgtccgggggtggacggaccggcggacgtccagggcgacgcggcggacccgggcagcggtagactagcttaaggtgtgcagagtccctgcggcaggcgacgaacacgaccgtcgtcgattagtaccggtacggtcggtggtatcgaagtcttgatcactgtacactagaggtaatacggttatccacagaatcaggggataacgcaggaaagaacatgtgagcaaaaggccagcaaaaggccaggaaccgtaaaaaggccgcgttgctggcgtttttccataggctccgcccccctgacgagcatcacaaaaatcgacgctcaagtcagaggtggcgaaacccgacaggactataaagataccaggcgtttccccctggaagctccctcgtgcgctctcctgttccgaccctgccgcttaccggatacctgtccgcctttctcccttcgggaagcgtggcgctttctcatagctcacgctgtaggtatctcagttcggtgtaggtcgttcgctccaagctgggctgtgtgcacgaaccccccgttcagcccgaccgctgcgccttatccggtaactatcgtcttgagtccaacccggtaagacacgacttatcgccactggcagcagccactggtaacaggattagcagagcgaggtatgtaggcggtgctacagagttcttgaagtggtggcctaactacggctacactagaaggacagtatttggtatctgcgctctgctgaagccagttaccttcggaaaaagagttggtagctcttgatccggcaaacaaaccaccgctggtagcggtggttttttt |
| pDK46  ARAC PROTEIN  GAM PROTEIN  BET PROTEIN  EXONUCLEASE  REPA101 PROTEIN  BETA LACTAMSE | catcgatttaTTATGACAACTTGACGGCTACATCATTCACTTTTTCTTCACAACCGGCACGGAACTCGCTCGGGCTGGCCCCGGTGCATTTTTTAAATACCCGCGAGAAATAGAGTTGATCGTCAAAACCAACATTGCGACCGACGGTGGCGATAGGCATCCGGGTGGTGCTCAAAAGCAGCTTCGCCTGGCTGATACGTTGGTCCTCGCGCCAGCTTAAGACGCTAATCCCTAACTGCTGGCGGAAAAGATGTGACAGACGCGACGGCGACAAGCAAACATGCTGTGCGACGCTGGCGATATCAAAATTGCTGTCTGCCAGGTGATCGCTGATGTACTGACAAGCCTCGCGTACCCGATTATCCATCGGTGGATGGAGCGACTCGTTAATCGCTTCCATGCGCCGCAGTAACAATTGCTCAAGCAGATTTATCGCCAGCAGCTCCGAATAGCGCCCTTCCCCTTGCCCGGCGTTAATGATTTGCCCAAACAGGTCGCTGAAATGCGGCTGGTGCGCTTCATCCGGGCGAAAGAACCCCGTATTGGCAAATATTGACGGCCAGTTAAGCCATTCATGCCAGTAGGCGCGCGGACGAAAGTAAACCCACTGGTGATACCATTCGCGAGCCTCCGGATGACGACCGTAGTGATGAATCTCTCCTGGCGGGAACAGCAAAATATCACCCGGTCGGCAAACAAATTCTCGTCCCTGATTTTTCACCACCCCCTGACCGCGAATGGTGAGATTGAGAATATAACCTTTCATTCCCAGCGGTCGGTCGATAAAAAAATCGAGATAACCGTTGGCCTCAATCGGCGTTAAACCCGCCACCAGATGGGCATTAAACGAGTATCCCGGCAGCAGGGGATCATTTTGCGCTTCAGCCATacttttcatactcccgccattcagagaagaaaccaattgtccatattgcatcagacattgccgtcactgcgtcttttactggctcttctcgctaaccaaaccggtaaccccgcttattaaaagcattctgtaacaaagcgggaccaaagccatgacaaaaacgcgtaacaaaagtgtctataatcacggcagaaaagtccacattgattatttgcacggcgtcacactttgctatgccatagcatttttatccataagattagcggatcctacctgacgctttttatcgcaactctctactgtttctccatacccgtttttttgggaattcgagctctaaggaggttataaaaaATGGATATTAATACTGAAACTGAGATCAAGCAAAAGCATTCACTAACCCCCTTTCCTGTTTTCCTAATCAGCCCGGCATTTCGCGGGCGATATTTTCACAGCTATTTCAGGAGTTCAGCCATGAACGCTTATTACATTCAGGATCGTCTTGAGGCTCAGAGCTGGGCGCGTCACTACCAGCAGCTCGCCCGTGAAGAGAAAGAGGCAGAACTGGCAGACGACATGGAAAAAGGCCTGCCCCAGCACCTGTTTGAATCGCTATGCATCGATCATTTGCAACGCCACGGGGCCAGCAAAAAATCCATTACCCGTGCGTTTGATGACGATGTTGAGTTTCAGGAGCGCATGGCAGAACACATCCGGTACATGGTTGAAACCATTGCTCACCACCAGGTTGATATTGATTCAGAGGTATAAaacgaATGAGTACTGCACTCGCAACGCTGGCTGGGAAGCTGGCTGAACGTGTCGGCATGGATTCTGTCGACCCACAGGAACTGATCACCACTCTTCGCCAGACGGCATTTAAAGGTGATGCCAGCGATGCGCAGTTCATCGCATTACTGATCGTTGCCAACCAGTACGGCCTTAATCCGTGGACGAAAGAAATTTACGCCTTTCCTGATAAGCAGAATGGCATCGTTCCGGTGGTGGGCGTTGATGGCTGGTCCCGCATCATCAATGAAAACCAGCAGTTTGATGGCATGGACTTTGAGCAGGACAATGAATCCTGTACATGCCGGATTTACCGCAAGGACCGTAATCATCCGATCTGCGTTACCGAATGGATGGATGAATGCCGCCGCGAACCATTCAAAACTCGCGAAGGCAGAGAAATCACGGGGCCGTGGCAGTCGCATCCCAAACGGATGTTACGTCATAAAGCCATGATTCAGTGTGCCCGTCTGGCCTTCGGATTTGCTGGTATCTATGACAAGGATGAAGCCGAGCGCATTGTCGAAAATACTGCATACACTGCAGAACGTCAGCCGGAACGCGACATCACTCCGGTTAACGATGAAACCATGCAGGAGATTAACACTCTGCTGATCGCCCTGGATAAAACATGGGATGACGACTTATTGCCGCTCTGTTCCCAGATATTTCGCCGCGACATTCGTGCATCGTCAGAACTGACACAGGCCGAAGCAGTAAAAGCTCTTGGATTCCTGAAACAGAAAGCCGCAGAGCAGAAGGTGGCAGCATGACACCGGACATTATCCTGCAGCGTACCGGGATCGATGTGAGAGCTGTCGAACAGGGGGATGATGCGTGGCACAAATTACGGCTCGGCGTCATCACCGCTTCAGAAGTTCACAACGTGATAGCAAAACCCCGCTCCGGAAAGAAGTGGCCTGACATGAAAATGTCCTACTTCCACACCCTGCTTGCTGAGGTTTGCACCGGTGTGGCTCCGGAAGTTAACGCTAAAGCACTGGCCTGGGGAAAACAGTACGAGAACGACGCCAGAACCCTGTTTGAATTCACTTCCGGCGTGAATGTTACTGAATCCCCGATCATCTATCGCGACGAAAGTATGCGTACCGCCTGCTCTCCCGATGGTTTATGCAGTGACGGCAACGGCCTTGAACTGAAATGCCCGTTTACCTCCCGGGATTTCATGAAGTTCCGGCTCGGTGGTTTCGAGGCCATAAAGTCAGCTTACATGGCCCAGGTGCAGTACAGCATGTGGGTGACGCGAAAAAATGCCTGGTACTTTGCCAACTATGACCCGCGTATGAAGCGTGAAGGCCTGCATTATGTCGTGATTGAGCGGGATGAAAAGTACATGGCGAGTTTTGACGAGATCGTGCCGGAGTTCATCGAAAAAATGGACGAGGCACTGGCTGAAATTGGTTTTGTATTTGGGGAGCAATGGCGATGACgcatcctcacgataatatccgggtaggcgcaatcactttcgtctactccgttacaaagcgaggctgggtatttcccggcctttctgttatccgaaatccactgaaagcacagcggctggctgaggagataaataataaacgaggggctgtatgcacaaagcatcttctgttgagttaagaacgagtatcgagatggcacatagccttgctcaaattggaatcaggtttgtgccaataccagtagaaacagacgaagaatccatgggtatggacagttttccctttgatatgtaacggtgaacagttgttctacttttgtttgttagtcttgatgcttcactgatagatacaagagccataagaaccTCAGATCCTTCCGTATTTAGCCAGTATGTTCTCTAGTGTGGTTCGTTGTTTTTGCGTGAGCCATGAGAACGAACCATTGAGATCATACTTACTTTGCATGTCACTCAAAAATTTTGCCTCAAAACTGGTGAGCTGAATTTTTGCAGTTAAAGCATCGTGTAGTGTTTTTCTTAGTCCGTTACGTAGGTAGGAATCTGATGTAATGGTTGTTGGTATTTTGTCACCATTCATTTTTATCTGGTTGTTCTCAAGTTCGGTTACGAGATCCATTTGTCTATCTAGTTCAACTTGGAAAATCAACGTATCAGTCGGGCGGCCTCGCTTATCAACCACCAATTTCATATTGCTGTAAGTGTTTAAATCTTTACTTATTGGTTTCAAAACCCATTGGTTAAGCCTTTTAAACTCATGGTAGTTATTTTCAAGCATTAACATGAACTTAAATTCATCAAGGCTAATCTCTATATTTGCCTTGTGAGTTTTCTTTTGTGTTAGTTCTTTTAATAACCACTCATAAATCCTCATAGAGTATTTGTTTTCAAAAGACTTAACATGTTCCAGATTATATTTTATGAATTTTTTTAACTGGAAAAGATAAGGCAATATCTCTTCACTAAAAACTAATTCTAATTTTTCGCTTGAGAACTTGGCATAGTTTGTCCACTGGAAAATCTCAAAGCCTTTAACCAAAGGATTCCTGATTTCCACAGTTCTCGTCATCAGCTCTCTGGTTGCTTTAGCTAATACACCATAAGCATTTTCCCTACTGATGTTCATCATCTGAGCGTATTGGTTATAAGTGAACGATACCGTCCGTTCTTTCCTTGTAGGGTTTTCAATCGTGGGGTTGAGTAGTGCCACACAGCATAAAATTAGCTTGGTTTCATGCTCCGTTAAGTCATAGCGACTAATCGCTAGTTCATTTGCTTTGAAAACAACTAATTCAGACATacatctcaattggtctaggtgattttaatcactataccaattgagatgggctagtcaatgataattactagtccttttcctttgagttgtgggtatctgtaaattctgctagacctttgctggaaaacttgtaaattctgctagaccctctgtaaattccgctagacctttgtgtgttttttttgtttatattcaagtggttataatttatagaataaagaaagaataaaaaaagataaaaagaatagatcccagccctgtgtataactcactactttagtcagttccgcagtattacaaaaggatgtcgcaaacgctgtttgctcctctacaaaacagaccttaaaaccctaaaggcttaagtagcaccctcgcaagctcggttgcggccgcaatcgggcaaatcgctgaatattccttttgtctccgaccatcaggcacctgagtcgctgtctttttcgtgacattcagttcgctgcgctcacggctctggcagtgaatgggggtaaatggcactacaggcgccttttatggattcatgcaaggaaactacccataatacaagaaaagcccgtcacgggcttctcagggcgttttatggcgggtctgctatgtggtgctatctgactttttgctgttcagcagttcctgccctctgattttccagtctgaccacttcggattatcccgtgacaggtcattcagactggctaatgcacccagtaaggcagcggtatcatcaacggggtctgacgctcagtggaacgaaaactcacgttaagggattttggtcatgagattatcaaaaaggatcttcacctagatccttttaaattaaaaatgaagttttaaatcaatctaaagtatatatgagtaaacttggtctgacagTTACCAATGCTTAATCAGTGAGGCACCTATCTCAGCGATCTGTCTATTTCGTTCATCCATAGTTGCCTGACTCCCCGTCGTGTAGATAACTACGATACGGGAGGGCTTACCATCTGGCCCCAGTGCTGCAATGATACCGCGAGACCCACGCTCACCGGCTCCAGATTTATCAGCAATAAACCAGCCAGCCGGAAGGGCCGAGCGCAGAAGTGGTCCTGCAACTTTATCCGCCTCCATCCAGTCTATTAATTGTTGCCGGGAAGCTAGAGTAAGTAGTTCGCCAGTTAATAGTTTGCGCAACGTTGTTGCCATTGCTACAGGCATCGTGGTGTCACGCTCGTCGTTTGGTATGGCTTCATTCAGCTCCGGTTCCCAACGATCAAGGCGAGTTACATGATCCCCCATGTTGTGCAAAAAAGCGGTTAGCTCCTTCGGTCCTCCGATCGTTGTCAGAAGTAAGTTGGCCGCAGTGTTATCACTCATGGTTATGGCAGCACTGCATAATTCTCTTACTGTCATGCCATCCGTAAGATGCTTTTCTGTGACTGGTGAGTACTCAACCAAGTCATTCTGAGAATAGTGTATGCGGCGACCGAGTTGCTCTTGCCCGGCGTCAATACGGGATAATACCGCGCCACATAGCAGAACTTTAAAAGTGCTCATCATTGGAAAACGTTCTTCGGGGCGAAAACTCTCAAGGATCTTACCGCTGTTGAGATCCAGTTCGATGTAACCCACTCGTGCACCCAACTGATCTTCAGCATCTTTTACTTTCACCAGCGTTTCTGGGTGAGCAAAAACAGGAAGGCAAAATGCCGCAAAAAAGGGAATAAGGGCGACACGGAAATGTTGAATACTCATactcttcctttttcaatattattgaagcatttatcagggttattgtctcatgagcggatacatatttgaatgtatttagaaaaataaacaaataggggttccgcgcacatttccccgaaaagtgccacctg |
| pET28a-WT-BM3  Kanamycin resistance  P450-BM3 Sequence  Origin of replication | ttgagatcctttttttctgcgcgtaatctgctgcttgcaaacaaaaaaaccaccgctaccagcggtggtttgtttgccggatcaagagctaccaactctttttccgaaggtaactggcttcagcagagcgcagataccaaatactgtccttctagtgtagccgtagttaggccaccacttcaagaactctgtagcaccgcctacatacctcgctctgctaatcctgttaccagtggctgctgccagtggcgataagtcgtgtcttaccgggttggactcaagacgatagttaccggataaggcgcagcggtcgggctgaacggggggttcgtgcacacagcccagcttggagcgaacgacctacaccgaactgagatacctacagcgtgagctatgagaaagcgccacgcttcccgaagggagaaaggcggacaggtatccggtaagcggcagggtcggaacaggagagcgcacgagggagcttccagggggaaacgcctggtatctttatagtcctgtcgggtttcgccacctctgacttgagcgtcgatttttgtgatgctcgtcaggggggcggagcctatggaaaaacgccagcaacgcggcctttttacggttcctggccttttgctggccttttgctcacatgttctttcctgcgttatcccctgattctgtggataaccgtattaccgcctttgagtgagctgataccgctcgccgcagccgaacgaccgagcgcagcgagtcagtgagcgaggaagcggaagagcgcctgatgcggtattttctccttacgcatctgtgcggtatttcacaccgcaatggtgcactctcagtacaatctgctctgatgccgcatagttaagccagtatacactccgctatcgctacgtgactgggtcatggctgcgccccgacacccgccaacacccgctgacgcgccctgacgggcttgtctgctcccggcatccgcttacagacaagctgtgaccgtctccgggagctgcatgtgtcagaggttttcaccgtcatcaccgaaacgcgcgaggcagctgcggtaaagctcatcagcgtggtcgtgaagcgattcacagatgtctgcctgttcatccgcgtccagctcgttgagtttctccagaagcgttaatgtctggcttctgataaagcgggccatgttaagggcggttttttcctgtttggtcactgatgcctccgtgtaagggggatttctgttcatgggggtaatgataccgatgaaacgagagaggatgctcacgatacgggttactgatgatgaacatgcccggttactggaacgttgtgagggtaaacaactggcggtatggatgcggcgggaccagagaaaaatcactcagggtcaatgccagcgcttcgttaatacagatgtaggtgttccacagggtagccagcagcatcctgcgatgcagatccggaacataatggtgcagggcgctgacttccgcgtttccagactttacgaaacacggaaaccgaagaccattcatgttgttgctcaggtcgcagacgttttgcagcagcagtcgcttcacgttcgctcgcgtatcggtgattcattctgctaaccagtaaggcaaccccgccagcctagccgggtcctcaacgacaggagcacgatcatgcgcacccgtggggccgccatgccggcgataatggcctgcttctcgccgaaacgtttggtggcgggaccagtgacgaaggcttgagcgagggcgtgcaagattccgaataccgcaagcgacaggccgatcatcgtcgcgctccagcgaaagcggtcctcgccgaaaatgacccagagcgctgccggcacctgtcctacgagttgcatgataaagaagacagtcataagtgcggcgacgatagtcatgccccgcgcccaccggaaggagctgactgggttgaaggctctcaagggcatcggtcgagatcccggtgcctaatgagtgagctaacttacattaattgcgttgcgctcactgcccgctttccagtcgggaaacctgtcgtgccagctgcattaatgaatcggccaacgcgcggggagaggcggtttgcgtattgggcgccagggtggtttttcttttcaccagtgagacgggcaacagctgattgcccttcaccgcctggccctgagagagttgcagcaagcggtccacgctggtttgccccagcaggcgaaaatcctgtttgatggtggttaacggcgggatataacatgagctgtcttcggtatcgtcgtatcccactaccgagatatccgcaccaacgcgcagcccggactcggtaatggcgcgcattgcgcccagcgccatctgatcgttggcaaccagcatcgcagtgggaacgatgccctcattcagcatttgcatggtttgttgaaaaccggacatggcactccagtcgccttcccgttccgctatcggctgaatttgattgcgagtgagatatttatgccagccagccagacgcagacgcgccgagacagaacttaatgggcccgctaacagcgcgatttgctggtgacccaatgcgaccagatgctccacgcccagtcgcgtaccgtcttcatgggagaaaataatactgttgatgggtgtctggtcagagacatcaagaaataacgccggaacattagtgcaggcagcttccacagcaatggcatcctggtcatccagcggatagttaatgatcagcccactgacgcgttgcgcgagaagattgtgcaccgccgctttacaggcttcgacgccgcttcgttctaccatcgacaccaccacgctggcacccagttgatcggcgcgagatttaatcgccgcgacaatttgcgacggcgcgtgcagggccagactggaggtggcaacgccaatcagcaacgactgtttgcccgccagttgttgtgccacgcggttgggaatgtaattcagctccgccatcgccgcttccactttttcccgcgttttcgcagaaacgtggctggcctggttcaccacgcgggaaacggtctgataagagacaccggcatactctgcgacatcgtataacgttactggtttcacattcaccaccctgaattgactctcttccgggcgctatcatgccataccgcgaaaggttttgcgccattcgatggtgtccgggatctcgacgctctcccttatgcgactcctgcattaggaagcagcccagtagtaggttgaggccgttgagcaccgccgccgcaaggaatggtgcatgcaaggagatggcgcccaacagtcccccggccacggggcctgccaccatacccacgccgaaacaagcgctcatgagcccgaagtggcgagcccgatcttccccatcggtgatgtcggcgatataggcgccagcaaccgcacctgtggcgccggtgatgccggccacgatgcgtccggcgtagaggatcgagatctcgatcccgcgaaattaatacgactcactataggggaattgtgagcggataacaattcccctctagaaataattttgtttaactttaagaaggagatataccatgggcagcagccatcatcatcatcatcacagcagcggcctggtgccgcgcggcagccatatggctagcatgactggtggacagcaaatgggtcgcATGACCATCAAGGAAATGCCACAGCCAAAGACTTTTGGGGAACTTAAGAACTTGCCCTTACTGAATACGGATAAGCCGGTCCAAGCACTTATGAAGATTGCAGATGAACTTGGTGAAATCTTCAAGTTCGAGGCCCCAGGACGCGTCACGCGTTATCTGAGCAGCCAACGTCTGATCAAAGAAGCTTGTGACGAGAGCCGCTTTGACAAGAATCTGTCGCAGGCATTGAAGTTCGTGCGCGATTTTGCGGGTGATGGATTGTTTACTTCATGGACTCACGAAAAAAATTGGAAAAAGGCCCATAATATCCTTTTACCTAGCTTTTCGCAACAGGCGATGAAAGGATACCACGCAATGATGGTCGATATCGCAGTGCAGTTGGTACAAAAATGGGAGCGTTTGAATGCTGACGAACACATCGAAGTCCCCGAAGATATGACGCGCTTGACGCTTGATACAATTGGGTTATGCGGCTTCAATTACCGCTTTAACTCTTTTTATCGCGATCAACCTCACCCATTTATTACATCGATGGTTCGTGCTTTGGATGAAGCGATGAATAAGCTTCAACGTGCAAATCCGGATGATCCTGCATATGACGAAAACAAGCGTCAGTTTCAAGAGGATATCAAGGTCATGAATGATTTAGTGGACAAAATTATCGCCGATCGTAAAGCATCGGGTGAACAATCTGACGATTTGCTTACACACATGCTGAACGGGAAAGACCCGGAGACCGGGGAGCCCCTTGACGATGAAAATATTCGTTACCAAATCATCACATTTTTAATTGCGGGCCATGAAACAACGTCAGGGCTTCTTTCATTTGCTTTATATTTTTTGGTTAAGAACCCGCATGTTTTACAGAAGGCGGCGGAGGAAGCCGCACGTGTTCTTGTTGACCCAGTACCATCATATAAGCAAGTCAAGCAACTGAAGTATGTAGGAATGGTGCTGAATGAGGCGCTTCGTCTGTGGCCCACGGCCCCCGCCTTCTCATTGTATGCTAAAGAGGACACAGTATTGGGAGGTGAATACCCTTTAGAGAAGGGGGACGAACTGATGGTCCTTATTCCACAGCTGCATCGCGACAAGACGATCTGGGGGGACGATGTAGAAGAATTCCGCCCTGAGCGTTTTGAAAATCCCAGTGCTATCCCTCAGCATGCATTTAAACCTTTCGGAAATGGACAACGTGCTTGCATTGGTCAACAATTTGCCCTTCATGAAGCCACCCTGGTTCTGGGAATGATGCTGAAACACTTCGACTTTGAGGACCACACTAATTATGAGTTAGATATTAAAGAGACCCTGACCTTGAAACCAGAGGGATTTGTGGTGAAAGCAAAGTCGAAAAAGATCCCCTTGGGTGGTATCCCCAGTCCAAGCACCGAGCAAAGCGCGAAAAAAGTCCGCAAAAAAGCTGAAAATGCCCACAATACTCCTCTTCTTGTATTGTACGGAAGCAATATGGGAACGGCGGAAGGAACTGCTCGTGATCTGGCGGACATCGCCATGAGTAAGGGATTCGCTCCCCAGGTTGCGACCTTGGATAGCCATGCCGGTAACTTGCCACGTGAGGGTGCGGTTTTGATCGTTACGGCCAGTTACAACGGTCACCCGCCGGACAACGCAAAGCAGTTTGTAGACTGGCTGGACCAAGCGTCTGCGGATGAGGTTAAAGGCGTTCGCTATTCGGTCTTTGGGTGTGGAGATAAGAACTGGGCGACTACTTACCAGAAGGTACCTGCGTTTATTGACGAAACCCTTGCCGCGAAGGGCGCTGAGAATATTGCGGATCGTGGCGAAGCGGATGCATCGGACGACTTCGAGGGTACGTACGAGGAATGGCGCGAACACATGTGGTCCGATGTGGCAGCATACTTTAACTTGGATATCGAGAACAGTGAGGATAATAAGAGCACCTTATCGCTTCAGTTCGTCGATAGTGCGGCCGATATGCCCTTGGCGAAAATGCACGGCGCATTTTCCACGAATGTAGTAGCTAGCAAAGAGTTACAGCAACCTGGGTCTGCCCGCTCCACCCGCCACCTTGAAATTGAACTTCCAAAAGAGGCGTCCTACCAAGAGGGTGACCATTTAGGGGTAATTCCGCGTAATTACGAGGGTATCGTTAATCGTGTGACCGCCCGTTTCGGGTTGGACGCTAGCCAACAAATTCGTTTGGAAGCAGAAGAGGAAAAGCTTGCGCACCTTCCTTTAGCGAAAACGGTGTCCGTGGAAGAGTTGCTGCAGTACGTAGAGTTGCAAGACCCAGTTACCCGCACCCAACTTCGCGCAATGGCTGCGAAGACGGTCTGTCCACCTCATAAAGTGGAGTTGGAAGCACTTCTTGAAAAGCAAGCATATAAGGAACAGGTCTTAGCGAAACGCTTGACCATGTTGGAACTTCTTGAGAAGTACCCTGCGTGCGAAATGAAGTTCAGCGAGTTTATTGCACTGCTTCCAAGTATTCGTCCCCGCTATTATTCAATCTCCAGCTCTCCGCGTGTCGATGAGAAGCAAGCCAGTATTACTGTCTCCGTGGTGTCAGGGGAAGCGTGGTCAGGGTACGGGGAATACAAAGGAATTGCTTCAAATTACCTTGCCGAGCTTCAGGAAGGGGATACGATTACTTGTTTCATTAGCACACCCCAATCAGAGTTTACGCTGCCAAAGGACCCCGAAACCCCTCTTATCATGGTAGGCCCAGGAACTGGGGTGGCCCCATTTCGTGGTTTTGTTCAAGCCCGTAAACAGCTTAAGGAACAGGGTCAATCCCTGGGTGAGGCGCACTTATACTTCGGGTGTCGCTCGCCTCACGAAGATTACTTATATCAGGAGGAGCTTGAGAACGCGCAATCTGAAGGAATCATTACATTGCACACAGCATTCAGCCGCATGCCAAATCAGCCCAAAACATATGTCCAGCACGTGATGGAGCAGGACGGTAAGAAGCTTATCGAATTGTTAGATCAGGGTGCTCATTTTTACATCTGCGGGGACGGCTCCCAAATGGCTCCGGCGGTGGAGGCAACGTTAATGAAAAGCTACGCGGATGTTCACCAGGTATCGGAGGCAGATGCGCGCTTATGGCTTCAACAGTTGGAGGAGAAGGGGCGTTACGCAAAAGACGTCTGGGCTGGGcaccaccaccaccaccactgagatccggctgctaacaaagcccgaaaggaagctgagttggctgctgccaccgctgagcaataactagcataaccccttggggcctctaaacgggtcttgaggggttttttgctgaaaggaggaactatatccggattggcgaatgggacgcgccctgtagcggcgcattaagcgcggcgggtgtggtggttacgcgcagcgtgaccgctacacttgccagcgccctagcgcccgctcctttcgctttcttcccttcctttctcgccacgttcgccggctttccccgtcaagctctaaatcgggggctccctttagggttccgatttagtgctttacggcacctcgaccccaaaaaacttgattagggtgatggttcacgtagtgggccatcgccctgatagacggtttttcgccctttgacgttggagtccacgttctttaatagtggactcttgttccaaactggaacaacactcaaccctatctcggtctattcttttgatttataagggattttgccgatttcggcctattggttaaaaaatgagctgatttaacaaaaatttaacgcgaattttaacaaaatattaacgcttacaatttaggtggcacttttcggggaaatgtgcgcggaacccctatttgtttatttttctaaatacattcaaatatgtatccgctcatgaattaattcttagaaaaactcatcgagcatcaaatgaaactgcaatttattcatatcaggattatcaataccatatttttgaaaaagccgtttctgtaatgaaggagaaaactcaccgaggcagttccataggatggcaagatcctggtatcggtctgcgattccgactcgtccaacatcaatacaacctattaatttcccctcgtcaaaaataaggttatcaagtgagaaatcaccatgagtgacgactgaatccggtgagaatggcaaaagtttatgcatttctttccagacttgttcaacaggccagccattacgctcgtcatcaaaatcactcgcatcaaccaaaccgttattcattcgtgattgcgcctgagcgagacgaaatacgcgatcgctgttaaaaggacaattacaaacaggaatcgaatgcaaccggcgcaggaacactgccagcgcatcaacaatattttcacctgaatcaggatattcttctaatacctggaatgctgttttcccggggatcgcagtggtgagtaaccatgcatcatcaggagtacggataaaatgcttgatggtcggaagaggcataaattccgtcagccagtttagtctgaccatctcatctgtaacatcattggcaacgctacctttgccatgtttcagaaacaactctggcgcatcgggcttcccatacaatcgatagattgtcgcacctgattgcccgacattatcgcgagcccatttatacccatataaatcagcatccatgttggaatttaatcgcggcctagagcaagacgtttcccgttgaatatggctcataacaccccttgtattactgtttatgtaagcagacagttttattgttcatgaccaaaatcccttaacgtgagttttcgttccactgagcgtcagaccccgtagaaaagatcaaaggatcttc |

**Table S7**: Concentration ranges for dose response curves

| Dose Response Curve Values (µM) | | | | |
| --- | --- | --- | --- | --- |
|  | Valencene (**1)** | Nootkatone (**2**) | Artemisinin (**3**) | Cedrol (**4**) |
| a | 0 | 0 | 0 | 0 |
| b | 125 | 12.5 | 1 | 12.5 |
| c | 250 | 25 | 4 | 25 |
| d | 500 | 50 | 11 | 50 |
| e | 1,000 | 100 | 33 | 100 |
| f | 2,000 | 150 | 100 | 150 |
| g | 2,500 | 200 | 300 | 200 |
| h | 3,000 | 250 | 500 | 250 |

**Table S8**: List of oligonucleotides used in this study

| **Name** | **Forward Primer**  **5’-3’** | **Reverse Primer**  **5’-3’** | **Purpose** |
| --- | --- | --- | --- |
| P1 | catagtgctgaaacgttgcggtcc | ttggcggcctcgagt | To revert MphR-13 residue 21 |
| P2 | gttgctcctgcattcggtcatcgctggc | tcagctgccgcagga | To revert MphR-13residue 147 |
| P3 | cgaggccgccaccatagtgctg | agtacctcgtcatcggacttgagc | To revert MphR-13residue 17 |
| P4 | cgaggccgccaccatagtgctgaaacgttgcggtcc | agtacctcgtcatcggacttgagc | To revert MphR-13residue 17 and 21. |
| P5 | catttaggtgggctaacatttgtttaactttaagaaggagatatacatatggctagc | ccaagctggagaccgaattcattatttgtagagctcatccatg | To amplify the αGFP insert for Gibson assembly |
| P6 | tgttagcccacctaaatgtaacagtcac | cggtctccagcttggctg | To amplify M7 to swap out *sf*GFP for αGFP |
| P7 | tttgatcgcgctaaatactgcttcaccacaaggaatgcaaacagtaatacaAggggtgtt | tgccttacgttcagacggggccgaagccccgtcgtcgtcattagaaaaactcatcgagca | To amplify the *kanR* gene from pET28a for the *tolC* deletion |

**Figure S1**: Surface areas of ErA and ligands **1**-**4**

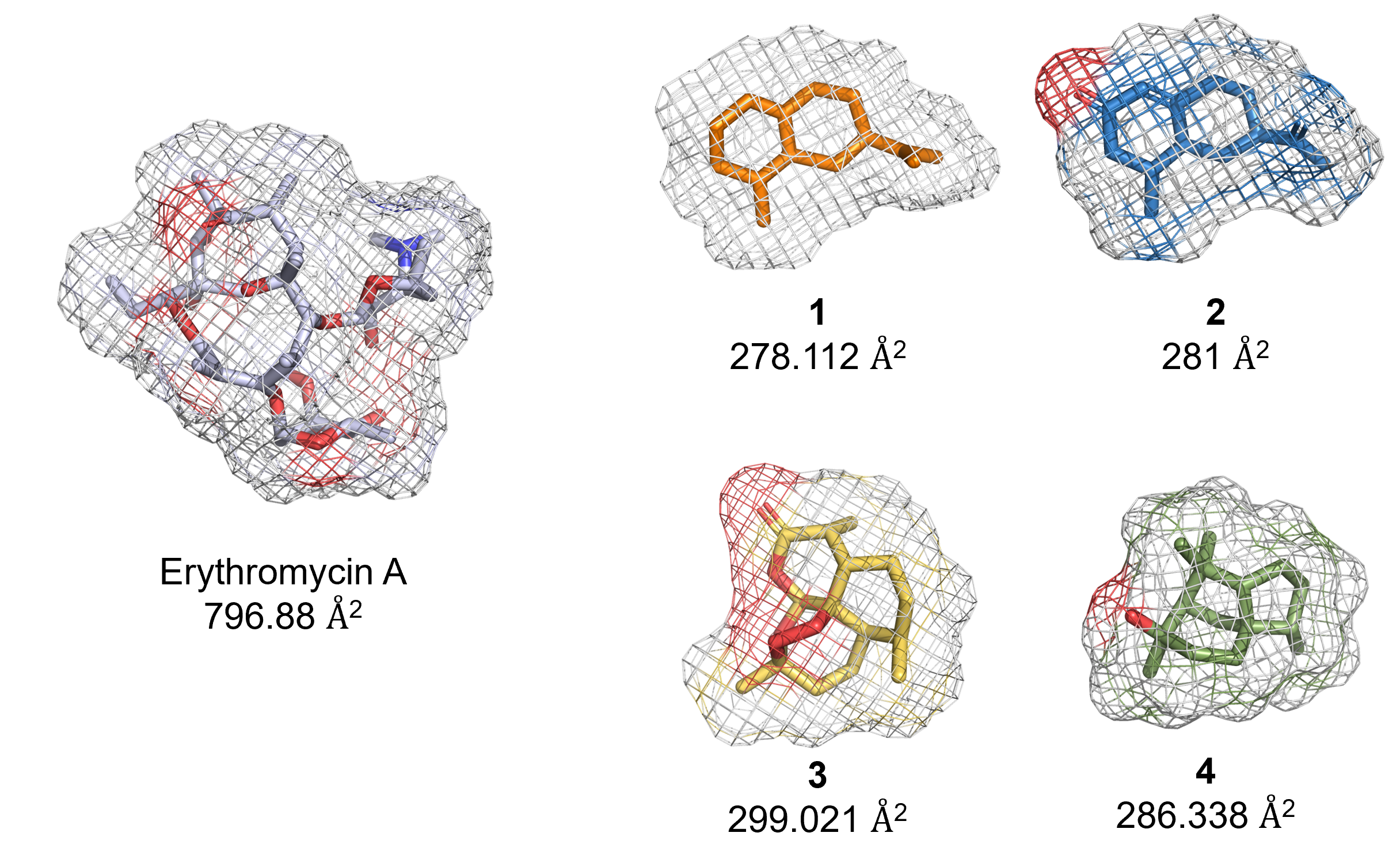
